# Using fluorescence flow cytometry data for single-cell gene expression analysis in bacteria

**DOI:** 10.1101/793976

**Authors:** Luca Galbusera, Gwendoline Bellement-Theroue, Arantxa Urchueguia, Thomas Julou, Erik van Nimwegen

## Abstract

Fluorescence flow cytometry is a highly attractive technology for quantifying single-cell expression distributions in bacteria in high-throughput. However, so far there has been no systematic investigation of the best practices for quantitative analysis of such data, what systematic biases exist, and what accuracy and sensitivity can be obtained. We here investigate these issues by systematically comparing flow cytometry measurements of fluorescent reporters in *E. coli* with measurements of the same strains in microscopic setups and develop a method for rigorous quantitative analysis of fluorescence flow cytometry data.

We find that forward and side scatter cannot be used to reliably estimate cell size in bacteria. Second, we show that cytometry measurements contain a large shot noise component that can be easily mistaken for intrinsic noise in gene expression, and show how calibration measurements can be used to correct for this measurement shot noise.

To aid other researchers with quantitative analysis of flow cytometry expression data in bacteria, we distribute *E-Flow*, an open-source R package that implements our methods for filtering cells based on forward and side scatter, and for estimating true biological expression means and variances from the fluorescence signal. The package is available at https://github.com/vanNimwegenLab/E-Flow.

## Introduction

In recent years, high-throughput measurements have played an increasingly important role in the advancement of biological research. The study of gene expression has been an area that has taken much advantage of these new technologies, and has reached the point where it is now possible to have genome-wide gene expression measurements at the single cell level for entire populations of cells. However, these high-throughput methods require accurate data analysis pipelines in order to quantify the biological signals and remove artifacts introduced by the measurement device.

Flow cytometry was initially developed in the 1940s for the identification of airbone bacteria and spores and thanks to the advancements of analytical cytology during the 1950s it became a more quantitative technology, allowing to determine and quantify the content of nucleic acids and proteins in living cells [1]. Today, sophisticated instruments are available that measure multiple optical and fluorescent characteristics of single cells. These have been extensively used both in the research and in the clinics, for example in immunology to distinguish cell types in blood samples. Flow cytometry is also a highly powerful technology for quantifying single-cell expression distributions in bacteria in high-throughput. Briefly, a beam of light is used to illuminate cells that flow one by one through a channel; a series of detectors is able to record the light scattered by the single cells at right angles or in the forward direction and the cell fluorescence stimulated by the incident light beam. Typically, each cell creates a signal of a given intensity and duration at the detectors, which the instrument reports as an “event”, summarizing it with three parameters, namely the height, the area and the width of the signal [2] [3].

Bacterial cells represent a challenge for the quantitative analysis and interpretation of the signals returned by the flow cytometer, due to their relatively small size and the low copy number of proteins and the consequently low fluorescence signal. All the more so as flow cytometers commonly available are optimized to measure eukaryotic cells. Since estimating concentrations of fluorescent reporters requires estimation of the cell volume, it is important to understand to which degree of accuracy the size of the single cells can be assessed from the scattering of the light, and how well viable cells can be separated from the debris. In addition, with regard to the estimation of fluorescence, it is important to be able to know to what extent variation in measured fluorescence intensity derives from biological variation, and to what extent it derives from measurement noise.

Here we rigorously investigate these questions using a systematic comparison of flow cytometry measurements with measurements from a microscopy setup. We find that, for rod-shaped bacteria, the relationship between cell size and forward- and side-scatter is too noisy to allow meaningful measurement of cell size variation. However, we also show that forward- and side-scatter can be used to distinguish true cells from debris and present a Bayesian mixture model to do so. For fluorescence measurements, we show that flow cytometry measurements contain a significant amount of shot-noise which can be easily mistaken for true biological variability, and develop a method to correct for this shot-noise using measurements of reference beads that are commonly used to calibrate flow cytometers. Using another mixture modeling approach we develop a rigorous method for estimating the true mean and variance in expression levels from fluorescence measurements, correcting for both autofluorescence and variance due to shot-noise. All these methods have been implemented as an R package called *E-Flow*, which can be easily integrated in any flow cytometry data analysis pipeline.

## Materials and methods

### Strains and growth conditions

We measured the fluorescence distributions for a number of different *Escherichia coli* MG1655 strains expressing cytoplasmic GFP from different promoters (corresponding to different mean expressions and noise levels), using both flow cytometry to measure batch cultures and time lapse microscopy to measure cells growing in a Mother Machine microfluidic device.

First, in previous work, we measured 24 hour long Mother Machine experiments with cells carrying a lacZ-GFP fusion and growing in M9 minimal media supplemented with 0.2% lactose (which leads to full induction of the lac operon), taking measurements every 3 minutes. A detailed experimental procedure is available elsewhere [4]. To obtain comparable measurements in flow cytometry, the same strain was grown overnight in M9 + 0.2% lactose, diluted 100× to fresh media and measured in mid-exponential phase (after approx. 4h), which is as close as possible to the conditions in the microfluidic device. Both in microfluidic experiments and in FCM experiments, the parent MG1655 strain without fluorescent reporter was measured in parallel to estimate the autofluorescence.

In addition, we measured fluorescence in single cells for a set of *E. coli* strains that carry a transcriptional reporter expressed from a low copy number plasmid [5], and growing in M9 + 0.4% glucose. These reporters are either for promoters regulated by lexA only (dinB, ftsK, lexA, polB, recA, recN, ruvA, or uvrD) [6] or for synthetic promoters obtained by experimental evolution so as to express at levels corresponding to the median or to 97^*th*^ percentile of all native *E. coli* promoters [7]. Throughout the paper, we refer to these two synthetic promoters as high and medium expressers. Mother Machine experiments were performed as described above [4], using M9 + 0.4% glucose (supplemented with 50*µg* / mL of kanamycin during the overnight preculture only), and acquiring data during 4 hours. For FCM measurements, samples were inoculated from glycerol stocks and grown overnight in 200*µL* of M9 + 0.4% glucose supplemented with 50*µg*/mL of kanamycin. Following a 100× dilution, cells were grown overnight again (without kanamycin), and finally diluted 100× to fresh medium. FCM measurements were recorded after 4h of growth (adjusting cells concentration with PBS if necessary). The strains used to measure the autofluorescence carried plasmids where GFP is not under the control of any promoter (puA66 and puA139) [5] and hence practically not expressed [7].

All cultures used for FCM measurements were incubated in 96-well plates at 37 °C with shaking at 600-650 rpm.

### Flow cytometry

The flow cytometry measurements were obtained with a BD FACSCanto II cytometer and were managed using the Diva 8 software. The excitation beam for the GFP was set at 488 nm and the emission signal was captured with a 530*/*30 nm bandpass filter. The gain voltage were set by default to 625V, 420V, and 600V for FSC, SSC, and GFP acquisition respectively, and events were created for measurements where *FSC >* 200 & *SSC >* 200. For each sample, 50000 events were recorded at a typical flow rate ranging from 10000 to 20000 per second.

### Calibration beads

CS&T (Cytometer Setup and Tracking Beads) are artificial fluorescent beads that are used to calibrate fluorescence measurement values [8]. We here used beads of lot 41720 that contains beads of two different sizes, which have high, medium (3*µm* in size) and low fluorescence (2*µm* in size) levels to characterize the shot noise introduced by the fluorescence measurement of the machine.

### R package *E-Flow*

The analysis pipeline presented in this paper has been implemented in the R package *E-Flow* available on GitHub https://github.com/vanNimwegenLab/E-Flow. Here the methods were tested with flow cytometers manufactured by BD and operated through the DIVA software. Nonetheless we kept the methods as general as possible, such that they should be applicable to flow cytometers of other manufacturers.

For a detailed explanation of the package, we refer to the GitHub page, to the vignette and to the documentation of the individual functions. Here we list the main components:

1. *Filtering*: The cells are filtered based on their scattering profile and an estimate of the mean and variance of the population is obtained. This is the most resource-intensive step and therefore can be parallelized.
2. *Mean and variance*: The mean and variance of the population of cells is computed. Measurements that are outliers in the fluorescence are accounted for using a mixture model.
3. *Autofluorescence removal*: Using the fluorescence distribution of non-expressing cells, an estimate of the autofluorescence is obtained and subtracted from the mean and variance of the population.
4. *Shot noise removal*: The shot noise introduced by the machine is removed and a corrected variance is calculated. This can be regarded as a proxy for the biological gene expression noise.

## Results

### Signals reported by the cytometer

Most flow cytometers, including the BD Canto II used here, report for each measured ‘event’ (typically corresponding to a single measured cell) a forward-scatter signal, a side-scatter signal, and a fluorescence signal. Each of these signals is in turn represented by 3 statistics of the electrical impulse, namely height, area, and width of the impulse (Fig 1). The height corresponds to the maximal value of the impulse, the area to the area under the curve and width is its time duration [2] (see Supplementary Material section 1.1.1). We noticed that these statistics are not all independent. In particular, for all three signals, the area is always directly proportional to the product of height and width (Suppl. Fig. S2 and Supplementary Material section 1.1.2). Moreover, while height and width vary approximately independently across events, the area correlates significantly with both (Suppl. Fig. S2). Therefore, we only use height and width for the subsequent analysis of the forward- and side-scatter signals. For the fluorescence signal, we were unable to find any systematic dependence between the width of the fluorescence signal and any biological signal, such as cell size or total fluorescence. In addition, for the calibration beads there is clearly no information in the width of the fluorescence signal (Suppl. Fig. S3). Therefore, for the fluorescence signal we will only use the height statistic as a proxy for the total fluorescence of the cells. While we believe that all these considerations apply generally to flow cytometers, we also observed anomalous behavior of the signal at very low fluorescence levels that may be specific to the BD machine used here (see Suppl. materials section 1.1.4). Due to this anomalous behavior, quantitative analysis is restricted to constructs for which the GFP fluorescence is at least as high as the autofluorescence of the cells (see Suppl. Fig. S4).

**Fig 1.**
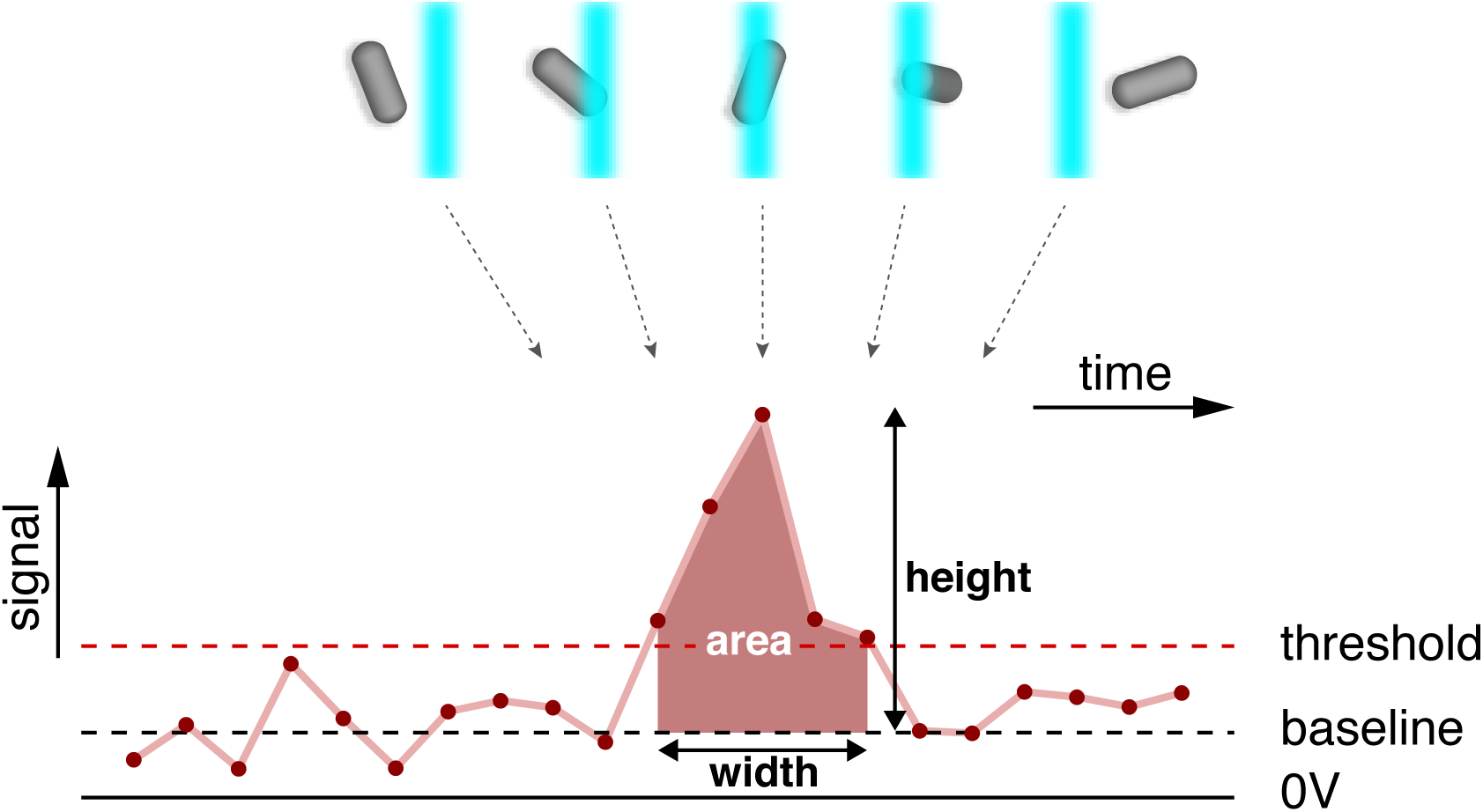
The signals reported by the cytometer. As a particle enters the laser beam, an electric signal (pulse) is generated which reaches its maximum when the particle is in the middle of the beam and trails off as the particle leaves the beam. Each pulse with height over a certain threshold is recorded and three quantities are reported: height, area, and width of the pulse.

### Filtering events based on their forward- and side-scatter

Bacterial cell don’t produce strong scattering signals in comparison to eukaryotic cells, we used a permissive setting of the device to call an event. This increases the likelihood of having spurious observations that correspond to non-viable cells and other debris. Consequently, we need a strategy for using the measured forward- and side-scatter of the events to separate viable cell measurements from debris. As explained above, the scatter of each event is characterized by 4 statistics, namely the height and width of both the forward- and side-scatter. Thus, the measured scatter of each event is represented by a point in a 4-dimensional space, and a given dataset corresponds to a distribution of points in this 4-dimensional space. To separate viable cells from debris we fit this distribution with a mixture of a multivariate Gaussian distribution and a uniform distribution, as detailed in the supplementary materials, section 1.2. The rationale behind this mixture modeling is that most of the data represents good cells and should cluster in this 4-dimensional space, whereas the outliers are relativel rare and more widely distributed. In this model, the Gaussian part of the mixture captures the cluster of good cells, while the uniform component takes care of outliers, i.e. fragments of dead cells and other debris.

Figure 2 shows the 4D scatter of forward- and side-scatter for events taken from *E. coli* cells that carry a lacZ-GFP fusion (see [4] for a description of the strain used) while growing in M9 minimal media supplemented with lactose. Besides the scatter of measurements, Fig. 2 also shows the multivariate Gaussian fitted to the data, showing that this Gaussian indeed captures the bulk of the measured events.

**Fig 2.**
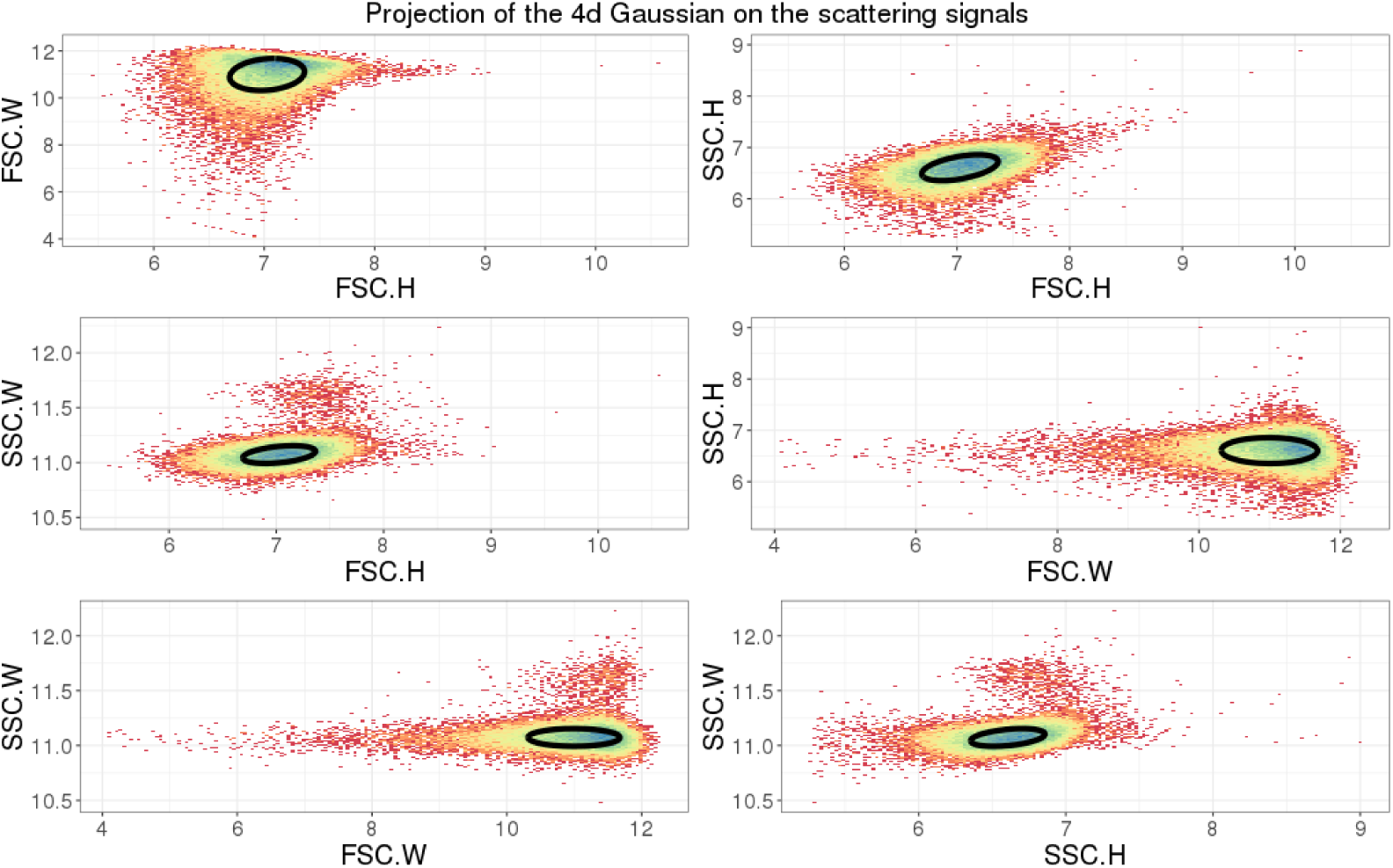
Mixture model fitting of the scatter signals. The panels show different two-dimensional projections of the full 4D distribution of forward- and side-scatter measurements for 5 × 10^4^ events obtained from *E. coli* cells growing in M9 minimal media with lactose. The ellipse shows the contour of the fitted multivariate Gaussian distribution, one standard deviation away in each principal direction. Note that the color indicates the local density of points.

Once the mixture model has been fitted to a dataset, a posterior probability *p*_*i*_ is calculated for each measured event *i* to correspond to a viable cell, i.e. that the observation derives from the multivariate Gaussian component of the mixture as opposed to deriving from the uniform distribution. By default the *E-flow* software retains all events with posterior probability *p*_*i*_ ≥ 0.5 and discards as outliers events with *p*_*i*_ < 0.5, but the user can change this threshold probability if desired. Suppl. Fig. S5 shows the same scatter of measured events as shown in Fig. 2, but now with selected events in red and events that were filtered out in black when using the default threshold of *p* = 0.5.

As the forward- and side-scatter should reflect the size, shape and composition of the objects measured in each event, one may wonder to what extent filtering out events based on their forward- and side-scatter may bias measurements towards cells of a certain size. Indeed, in previous work, e.g. [9], researchers have attempted to select subsets of cells with similar shapes and size by very strictly gating on forward- and side-scatter, retaining only those cells that lie near the center of the Gaussian distribution. To check the viability of such an approach, we compared the distribution of measured fluorescence levels with two extreme filtering strategies: one very lenient in which all events with *p > e*^−10^ are retained and one very strict in which only cells with *p* > 1 − *e*^−10^ are retained. As shown in Suppl. Fig. S6, there is virtually no difference in the observed distribution of fluorescence levels between the very lenient and very strict filtering. Given that we expect total fluorescence to scale with cell size, this observation suggests that strict filtering on forward- and side-scatter is not effective for selecting out a subset of cells with similar size.

### Forward- and side-scatter are poor measures of bacterial cell size

The extent to which forward- and side-scatter measurements of FCMs can be used to estimate the size of the measured object is a topic of much debate in the literature. It is generally assumed that forward scatter mostly reflects cell size, and that side scatter reflects surface properties such as granularity [1]. Here we used a comparison of identical cell populations in microscopy and FCM measurements to characterize the accuracy of forward- and side-scatter as a measure of the size of bacterial cells. In previous work we have established that time-lapse microscopy measurements of cells growing in microfluidic devices can measure cell size with an accuracy of around 3% error and GFP copy-number *G* with an error of about 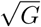 [4]. Using such microscopy measurements on *E. coli* cells carrying a lacZ-GFP fusion gene in its native locus while growing in M9 minimal media with lactose, we find a high correlation between lacZ-GFP levels and cell size (Fig. 3, top panels). That is, because lacZ-GFP concentrations fluctuate only moderately from cell to cell, and both size and GFP level measurements have high accuracy, the measured length captures around 70% of the variation in fluorescence measurements. However, as shown in the middle and bottom panels of Fig. 3, for the same cells grown in the same conditions in the FCM, neither the forward-nor side-scatter measurements show substantial correlation with fluorescence measurements.

**Fig 3.**
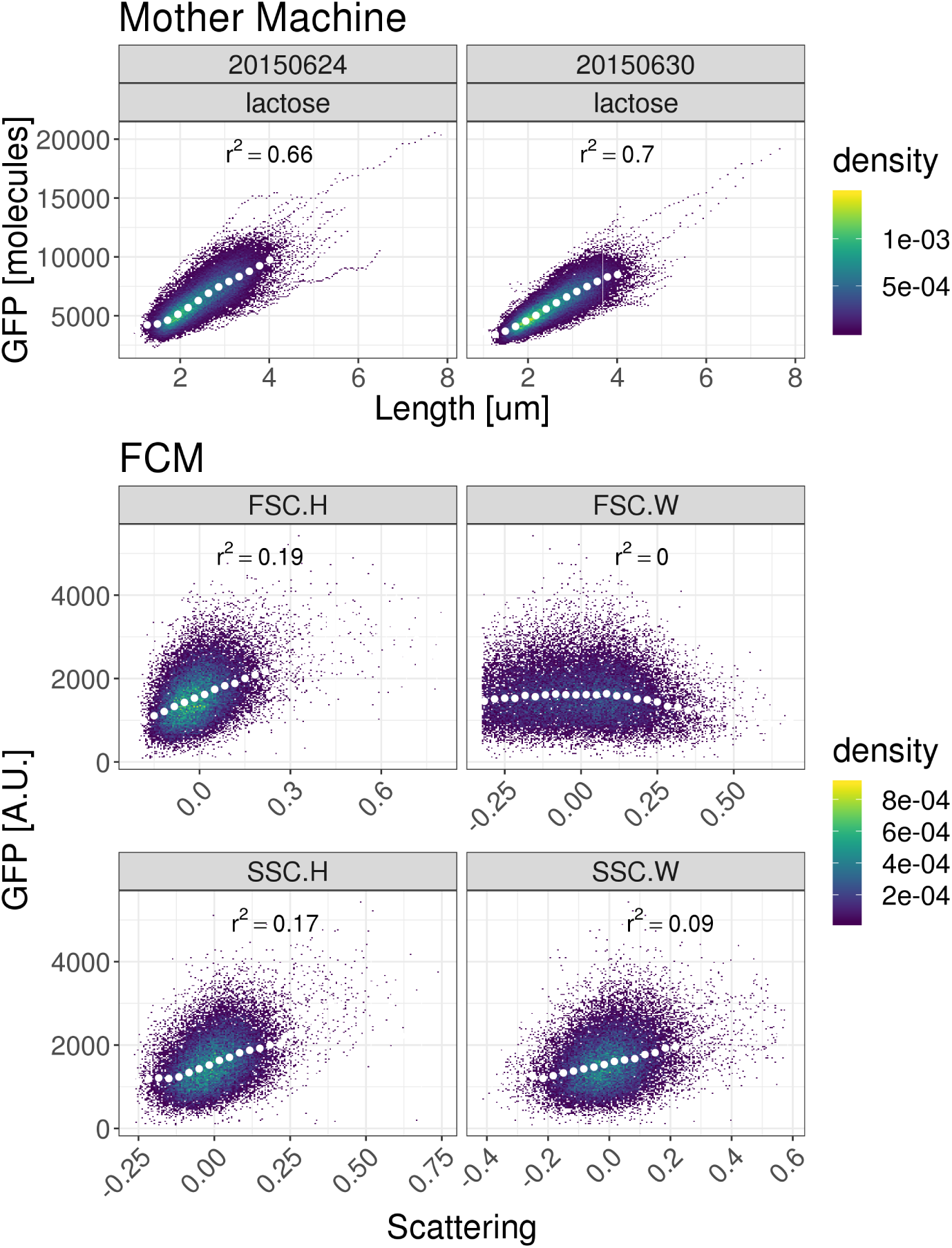
Correlation between cell size and fluorescence measurements for microscopy and FCM measurements. Each panel shows measured GFP fluorescence (vertical axis) and cell size estimates (horizontal axis) of cells growing in M9 minimal media with lactose. The top 2 panels show microscopy measurements from a microfluidic device [4]. The lower 4 panels show fluorescence measurements as a function of size estimates based on forward-(middle 2 panels) or side-scatter (bottom 2 panels) measurements in the FCM. The squared Pearson correlations between fluorescence and size measurements are indicated in each panel. Note that the color indicates the density of points. The white dots show median values of equally spaced bins along the horizontal axis.

The lack of correlations between size and fluorescence measurements in the FCM provides further evidence that the forward- and side-scatter do not provide accurate estimates of cell size for *E. coli*. To quantify how well forward- and side-scatter estimate cell size, we used the fact that the size of the errors of the microscopic measurements have been well characterized previously [4]. Since errors in size measurements in the microscope are proportional to cell size we work in log-space and denote by *x*_*t*_ the true log-size of a cell, by *x*_*m*_ its measured size and by *ϵ* the size of the error in the measurement. We thus have

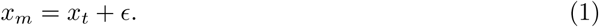

Because the error in the measurement *ϵ* is uncorrelated with the true value of the log-size *x*_*t*_, the variance of measured log-sizes *x*_*m*_ is also the sum of the variances in true log-size and measurement errors, i.e.

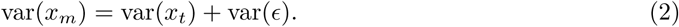

Since the variance of microscopy measurement errors is about var(*ϵ*) ≈ 0.03^2^, we can use the variance var(*x*_*m*_) of the measured log-sizes in the microscope to estimate the variance var(*x*_*t*_). Because equation (2) also applies to measurements in the FCM, and assuming that the variance var(*x*_*t*_) in true sizes is the same in the FCM, we can then finally use the measured variance var(*x*_*m*_) of size estimates in the FCM to estimate the variance var(*ϵ*) of the errors in size measurements in the FCM. Suppl. Fig. S7 shows the results of this analysis when using either the width or height of either the forward- or side-scatter measurements. The widths of the forward- and side-scatter signals carry virtually no information about cell size. While the heights of the forward- and side-scatter signals carry some information on cell size, the size of the measurement error is large in comparison to the true variability in cell sizes, such that these measurements cannot be used for effective estimates of cell size either. In short, our analysis shows that forward- and side-scatter measurements cannot be used to estimate variation in cell sizes of *E. coli*. Consequently, it will also be impossible to use the FCM measurements for estimating the GFP *concentration* of each cell. Thus, in the following we will focus on measuring the variation in *total* fluorescence across cells.

### Estimating the mean and variance of the fluorescence distribution

As has been observed by others [10], we observed that for virtually all *E. coli* promoters, the distribution of fluorescence levels is fitted very well by a log-normal distribution [7], i.e. the log-fluorescence follows a Gaussian distribution. Our *E-Flow* package fits a Gaussian distribution to the observed log-fluorescence levels, estimating a mean *µ* and variance *υ* for a given population of cells. We noticed that, even after filtering events on forward- and side-scatter as explained above, there are still clear outlying events, i.e. with fluorescence levels that lie far outside the range observed for almost all other events. To separate these outliers from valid measurements we modeled the distribution of log-fluorescence levels as a mixture of a Gaussian and a uniform distribution, fitting its parameters using expectation maximization (see section 1.3.1 of the Supplementary Material for details). The *E-Flow* package calculates an estimated mean *µ* and variance *υ* of the log-fluorescence levels of a set of measurements, together with error bars *σ*_*µ*_ and *σ*_*υ*_ on these estimates. In addition, transforming from log-fluorescence back to fluorescence in linear scale, the package also calculates mean and variance of the distribution of fluorescence levels, together with error bars (Suppl. Mat. 1.3.1).

### Autofluorescence estimation

It is well known that the laser used to excite the GFP can also excite other cellular components of the cell, resulting in an “autofluorescence” signal that also occurs in cells without GFP molecules. In addition, the fluorescence signal may also contain a background fluorescence component coming from sources other than the cell’s autofluorescence. In order to estimate GFP levels, we need to correct for these other sources of fluorescence and the *E-Flow* package allows for such correction by using measurements of cells that do not express GFP. Let’s call *I*_*M*_ the measured fluorescence intensity, *I*_*T*_ the true intensity (deriving from GFP molecules) and *A* the component from other sources of fluorescence, which for simplicity we will refer to as autofluorescence. We have the relation

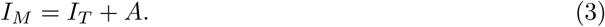

Assuming that the component *A* fluctuates independently from the true fluorescence *I*_*T*_, we obtain

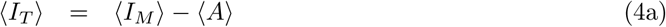

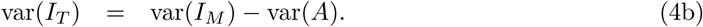

Using the procedure described in the previous section, we can estimate both 〈*A*〉 and var(*A*) from measurements of cells that either lack GFP, or where the GFP gene is known not to be expressed. Using these estimates, we can then infer the true mean and variance of GFP expression in cells carrying an active reporter using equation (4).

We measured autofluorescence levels *A* using strains carrying two different plasmids not expressing GFP, designed as negative control (see materials and methods) on 4 different days, measuring each strain in triplicate on each day. Figure 4 shows the estimated mean fluorescences (left 4 panels) and variances in fluorescences (right 4 panels) for each replicate of each strain (red an blue) on each day (one panel per day). Using a procedure described in Suppl. Mat. section 1.4, we averaged over different replicates on each day to calculate a mean fluorescence *µ*_*d*_ for each day (black line in each panel) and an error bar on this estimate (grey region in each panel), and similarly for the variances on each day (right 4 panels). We then additionally averaged over different days to calculate an overall average mean autofluorescence 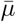 and an overall average variance in autofluorescence 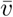 (see Suppl. Mat. 1.4).

**Fig 4.**
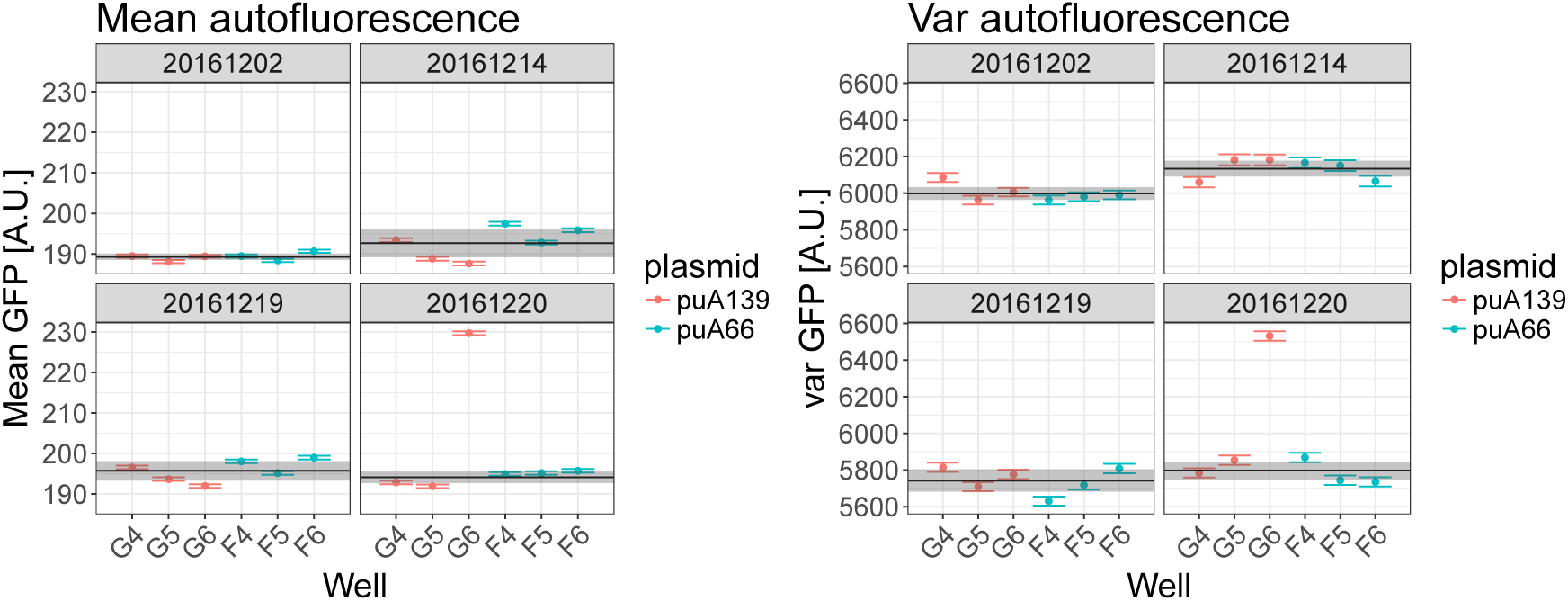
Autofluorescence measured on multiple days. Each panel shows the measured mean fluorescence (left 4 panels) and variance in fluorescence (right 4 panels) on a day, with each bar indicating the measured value and error bar for one replicate. Two different strains were used (indicated in red and blue) and each was measured in triplicate on each day. The black line and grey bar indicate the estimate day-dependent averages *µ*_*d*_ and their error-bars *σ*_*d*_. Note that well G6 on 20/12/2016 appears to be an outlier, possibly due to contamination of the well, which was excluded from the analysis.

### FCM fluorescence measurements exhibit significant shot noise

We used equation (4) to remove the autofluorescence contribution from the mean expression and variance of the population for a number of different transcriptional reporters and calculated the observed squared coefficient of variation *CV*^2^ for each promoter. Next, we took microscopy measurements from our microfluidic setup of the same *E. coli* strains growing in the same conditions, to measure *CV*^2^ for each of these promoters as well. As shown in the top panel of Fig. 5, we observe systematically higher *CV*^2^ in the FCM than in the microscopy setup and the difference in the two *CV*^2^s decreases almost exactly inversely with the mean expression level.

**Fig 5.**
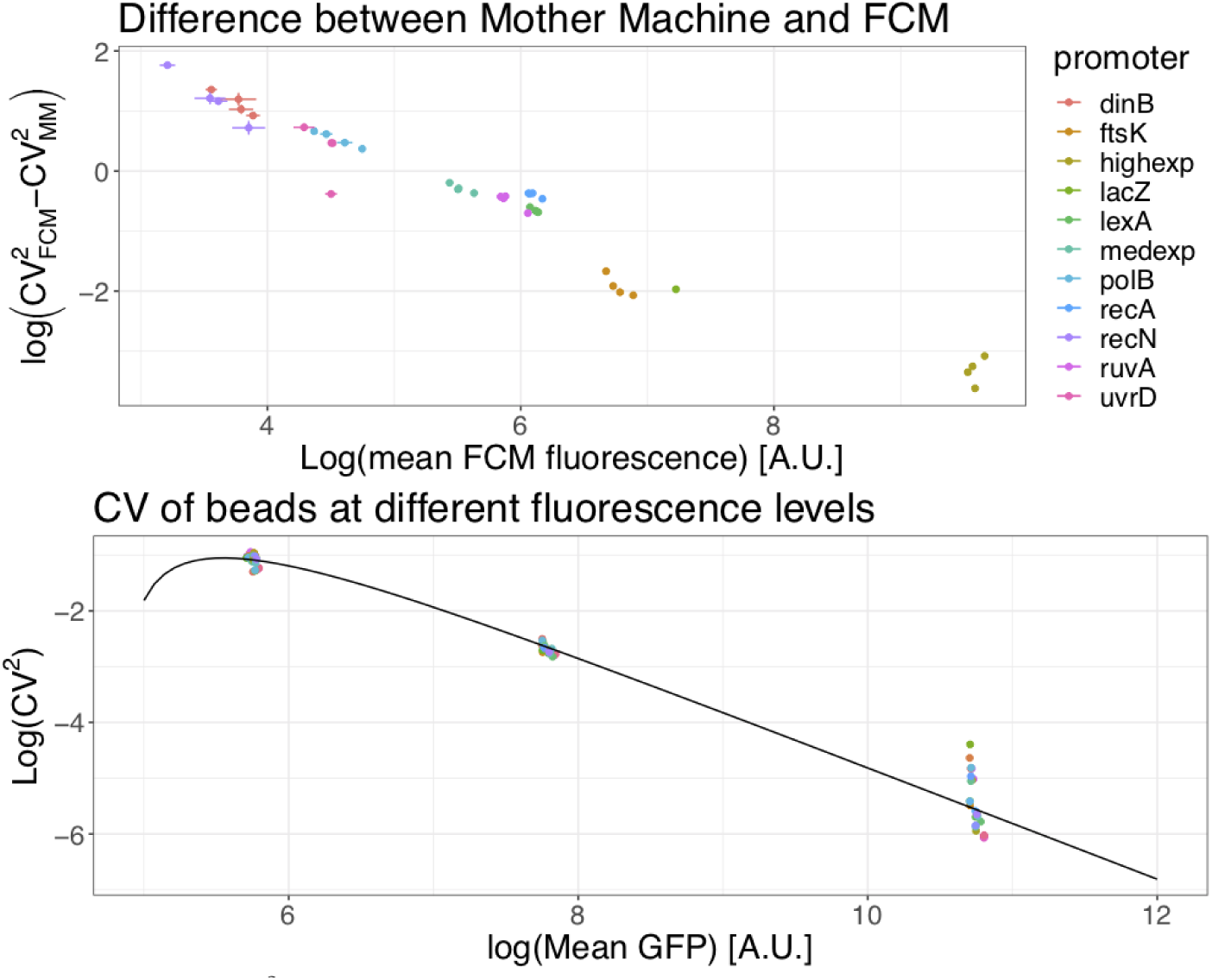
Difference in *CV*^2^ between the FCM and microscopy measurements shows FCM measurements contain substantial shot noise. *Top:* Difference between the *CV*^2^ as measured by the FCM and the microscopy setup for several different transcriptional reporters of *E. coli* promoters (colored points). Both axes are shown on a logarithmic scale. The difference in *CV*^2^ scales inversely with mean expression. *Bottom:* The observed *CV*^2^ of calibration beads of three different intensities also decreases as the inverse of mean intensity and this dependence can be well modeled by shot noise (black line), as given by equation (5).

Since the growth conditions in the FCM and the microfluidic setup were kept as close as possible, the true *CV*^2^ of the distribution of total GFP levels should be highly similar, so that the difference between the measured *CV*^2^ must derive from measurement noise. Indeed, one source of noise whose contribution to *CV*^2^ is expected to scale inversely with mean intensity is shot noise from the photomultiplier tube, whose *CV*^2^ scales as 1/mean [11]. Due to this noise, one generally has the following relationship between the measured fluorescence intensity *I*_*M*_ and the true intensity *I*_*T*_:

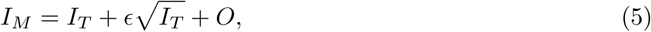

where *ϵ* is a Gaussian random variable with mean 0 and an (unknown) variance *δ*^2^ which quantifies the size of the shot noise. The constant term *O* is an offset that is added in BD devices in order to prevent the clipping of negative values during the digital conversion, when true intensities *I*_*T*_ are close to zero [2].

In order to estimate the size of the shot noise *δ*, we used a set of synthetic beads used to calibrate the FCM, which consists of a mixture of beads of three different fluorescent intensities. As shown in the bottom panel of Fig. 5 (and Suppl. Fig. S8) the *CV*^2^ of the artificial beads also drops inversely with mean expression. If we assume that the true variation of the beads can be ignored, we get from equation (5) that the measured *CV*^2^ is

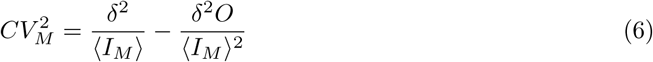

If we define 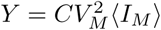 and 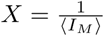, we obtain

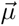

and we can infer both the strength *δ* and the offset *O* by fitting *Y* as a simple linear function of *X*. This simple approach leads to an inferred value of *δ* = 13.4 and *O* = 128. In the Supplementary Material section 1.5 we also present a more sophisticated Bayesian mixture model approach to inferring these quantities, which does not ignore the true variability of the beads, but assumes that the *CV*^2^ of the true intensities *I*_*T*_ is the same for all three types of beads. Using this more rigorous procedure, the resulting strength and offset are: *δ* = 12.7±0.6, *O* = 97±29 (Suppl. Fig. S8), which are close to the values we would have obtained with the more simple linear model of equation (6). Using this result we can now fit the observed *CV*^2^ that we expect to see; the fit describes well the observed data, as shown in the bottom panel of Fig. 5 (and in the top left panel of Suppl. Fig. S8).

Finally, we note that section 1.6.1 of the Supplementary Material covers two more subtle technical points that might affect the direct comparison of FCM measurements and microscopy measurement from growth in the the microfluidic device. First, one could argue that the age-distributions of the population of cells in the microfluidic device and in a population that is growing exponentially (i.e. as used in the FCM) are different. That is, since in the microfluidic device some newborn daughters are constantly washed out of the growth channels, there are relatively fewer cells close to birth and more cells close to division in the microfluidic device than in a population undergoing exponential growth in bulk (Suppl. Fig. S9). Since total fluorescence correlates with cell size, which again correlates well with time since birth, the access of ‘old’ cells could in principle effect the distribution of total fluorescence one observes. However, as shown in Suppl. Mat. section 1.6.1, we derive theoretically that the effects of the altered age-distribution are small enough to be neglected (Suppl. Fig. S10). Second, since in the microfluidic setup we measure the fluorescence of a cell multiple times during its cell cycle, there are clearly substantial correlations between different measurements and one might wonder whether this could significantly affect the observed statistics. In Suppl. Mat. section 1.6.2 we show that this effect is also negligible (Supplementary Fig S10).

### Correcting for autofluorescence and shot noise

After having estimated the mean and variance of the autofluorescence, and the strength of the FCM’s shot noise, we can now correct the measured means and variances of transcriptional reporters for these two components. Combining the autofluorescence contribution from equation (3) and the shot noise component from equation (5), we can write the measured intensity *I*_*M*_ as

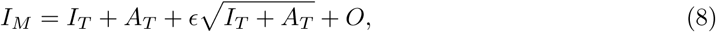

and the measured autofluorescence as

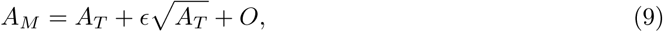

where variables with subscript *T* correspond to true values and variables with subscript *M* correspond to measured values, *ϵ* is again a Gaussian distributed variable with mean zero and variance *δ*^2^ and *O* is a constant offset. From these equations we find for the mean and variance of the measured intensities *I*_*M*_:

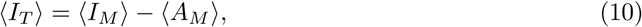

and

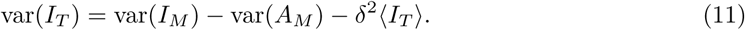

Using these expressions we calculated 〈*I*_*T*_〉, var(*I*_*T*_) and the resulting *CV*^2^ for a set of different *E. coli* promoters and compared the results with the *CV*^2^ measured for the same promoters in the microscopy setup. As shown in Fig. 6, while the estimates *CV*^2^ are still a little bit larger in the FCM, they are much closer to the results obtained with the microscopy measurements and the difference no longer systematically depends on the mean expression level.

**Fig 6.**
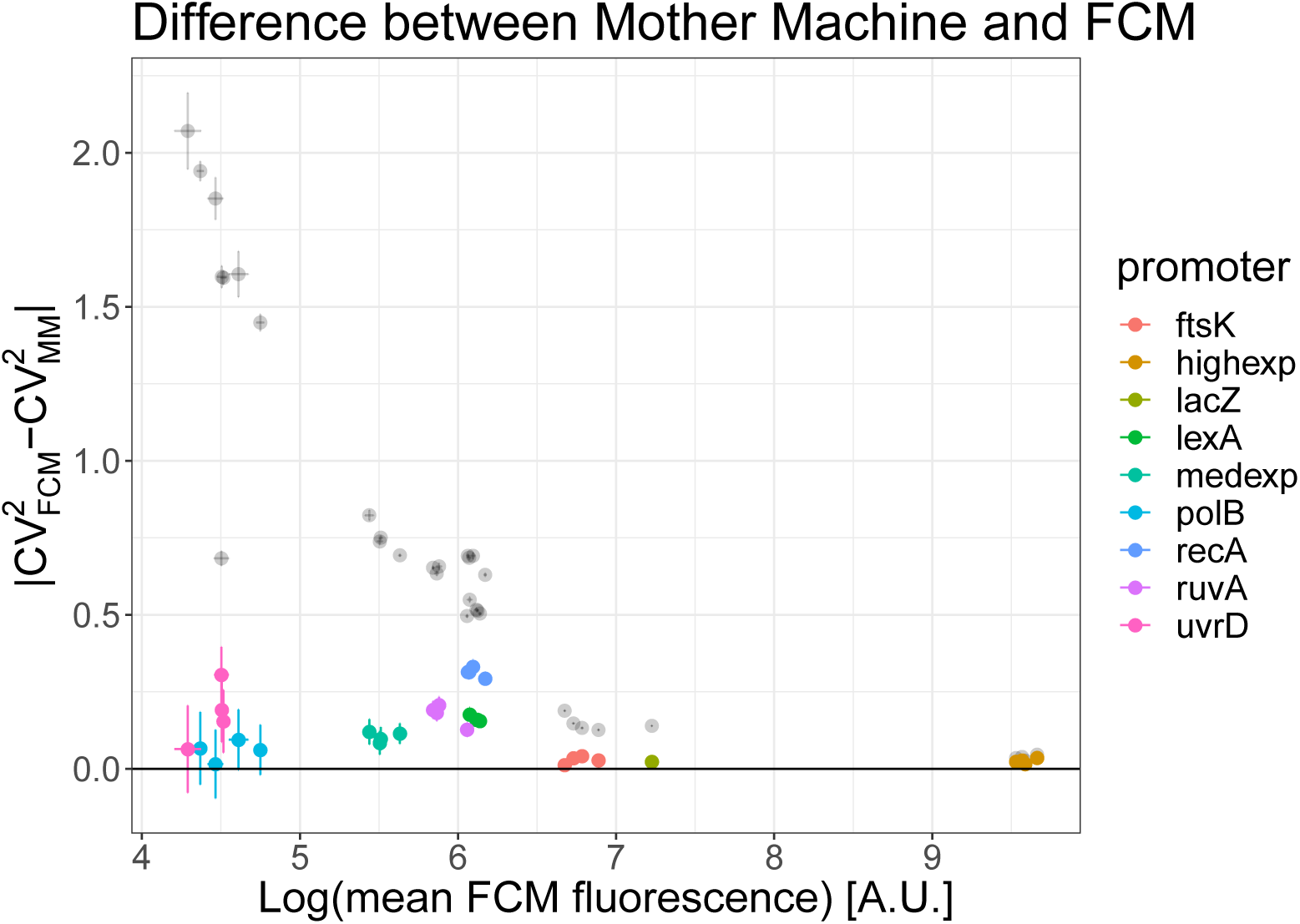
Comparison of *CV*^2^ from FCM and microscope measurements after correcting for autofluorescence and shot noise. Difference between the *CV*^2^ of different transcriptional reporters of native *E. coli* promoters between FCM and microscope measurements. The black transparent dots use uncorrected FCM measurements and reproduce Fig. 5 in linear scale, while the coloured dots are the *CV*^2^ after correcting for the shot noise. Only promoters expressing more than exp(4) above the autofluorescence were considered.

## Discussion

Although flow cytometry is an attractive technology for single-cell analysis of gene expression in high-throughput, we have shown that for data from bacterial cells there are a number of challenges in data analysis procedures to overcome in order to obtain accurate quantification. We here developed a number of procedures for measuring single-cell expression distributions in bacteria using FCM data and implemented them in an R package called *E-Flow*.

We first analyzed the forward- and side-scatter signals and their correlation structure. There seems to be little agreement in the literature as to when to use forward-scatter or side-scatter and whether to use height, width or area. We showed that only width and height provide independent measurements and developed a Bayesian mixture model for separating viable cell measurements from debris and other outliers using the full 4-dimensional distribution of forward- and side-scatter measurements. In general the filter we developed is much broader than the very strict gating strategies that are sometimes used and it discards only a small fraction of the events.

We next investigated to what extent forward- and side-scatter can be used to estimate the size of each measured cell. In the analysis of eukaryotic cells, the forward- and side-scatter are often used to separate different populations of cells characterized by different sizes and granularity, e.g. in the separation of different types of blood cells [12]. Nonetheless, these techniques are more qualitative than quantitative and it is debated how much precision one can achieve in relating the scattering signals to the size of the cells [1]. Here we have shown that for bacteria, although there is a correlation between cell size and forward- and side-scatter, this correlation is very weak and can in practice not be used to meaningfully estimate cell size. In particular, it is not possible to use either a strict gating strategy or estimation of cell size based on forward- and side-scatter, to estimate GFP concentrations of individual cells. Consequently, FCM measurements on bacterial cells can only be used to obtain distributions of total fluorescence per cell and not GFP concentrations.

We developed another Bayesian mixture model for accurately estimating the mean and variance of fluorescence levels for a population of cells. In addition, using cells that do not express fluorescent reporters, these techniques can also be used to estimate the distribution of autofluorescence, and we provide methods for removing the autofluorescence contribution from the observed means and variances of cells carrying fluorescent reporters. This step is fundamental since generally the autofluorescence cannot be ignored as just a small perturbation to the overall fluorescence signal.

Using data both from transcriptional reporters in *E. coli* as well as from artificial calibration beads, we showed that FCM fluorescence measurements contain a substantial amount of shot noise. This shot noise can be easily mistaken for biological variability in gene expression and it is thus crucial to correct for it in order to properly quantify gene expression noise. We developed a method that uses data from calibration beads to estimate the strength of this shot noise, and to correct for it in the estimation of means and variances of fluorescent reporters across a population of cells. We showed that, only after correcting for shot noise do gene expression noise measurements from the FCM converge to those obtained from microscopy measurements.

Although our data have been applied to flow cytometers manufactured by BD, we believe that the results are also valid for a broad range of other devices, as the autofluorescence is a phenomenon intrinsic to the cells, and the shot noise emerges in any device that uses a photomultiplier tube to transform a light signal into an electrical current.

## 1 Supplementary Material

### 1.1 Flow cytometers signals and their statistical properties

#### 1.1.1 Signal acquisition

In the flow cytometer, each cell is made to pass through a set of laser beams in order to measure its scattering profile and its fluorescence intensity at several wavelengths. The scatter has two components, forward- and side-scatter, which are usually interpreted as follows [3]:

1. Forward-scatter (FSC) generally reflects the size of the cells or particles.
2. Side-scatter (SSC) generally reflects the internal complexity or granularity of the cells or particles.

The scatter and fluorescence of each event (typically a cell) is converted by a photomultiplier tube into a pulse of electrical signal that is characterized by a height and a duration (Fig. 1). Roughly speaking, a pulse begins when a particle enters the laser beam and it reaches its maximum when the particle reaches the middle of the beam. In order to represent the pulse, the signal is sent to an analog-to-digital converter (ADC) that discretizes the continuous pulse into a digital one. For the machine used in this study, the ADC does this by sampling the pulse every 0.1 *µ*sec and the height of the signal is mapped to an integer between 0 and 2^14^, i.e. 14 bits are used to quantify the height of the signal [2].

#### 1.1.2 Correlations among the signals

The cytometer gives three statistics of a measured pulse (height, width, and area), and we investigated correlations between these statistics. For the machine used in our study, we find that the area is in fact exactly proportional to the product of the height and width of the signal (Suppl. Fig. S1).

Consequently, of the three statistics reported, only two are independent and we next measured the correlations between all pairs of statistics to determine which of these are most independent. As shown in Suppl. Fig. S2 we can see that area correlates significantly with both height and width whereas height and width do not show significant correlation. We thus decided to characterize each signal by height and width.

#### 1.1.3 Signal width is not informative for fluorescence measurements

To assess the relative importance of the height and width statistics for fluorescence measurements we made use of the calibration beads manufactured by BD, which come in a set of three different expression levels. Supplementary Fig. S3 shows that, whereas the heights of the signal pulses show three main peaks, corresponding to the three intensities of the beads, the distribution of measured widths is almost identical for all beads. Consequently, we chose to only use signal height to quantify fluorescence.

#### 1.1.4 Anomalous behavior at low fluorescence intensities

As can already be observed from the measurements of the calibration beads (Suppl. Fig. S3), due to shot noise in the fluorescence measurements, the coefficient of variation generally increases as the fluorescence level decreases. This general negative correlation between mean fluorescence and its coefficient of variation can not only be observed for the calibration beads, but also for a library of transcriptional reporters of native *E. coli* promoters (Suppl. Fig. S4). However, as Suppl. Fig. S4 also shows, when fluorescence levels become very low, the *CV*^2^ of the measurements reaches a maximum and starts to *decrease* as the mean decreases further, and the lowest *CV*^2^ is observed for cells without any GFP expression. We hypothesize that this decrease in *CV*^2^ for very low fluorescence levels derives from the fact that autofluorescence levels vary less than GFP fluorescence levels, in combination with some specific signal processing that is performed by the machine at low signal intensities. Although we invested a significant amount of time in trying to reverse engineer what signal processing may cause this anomalous behavior at very low fluorescence levels, these attempts were ultimately not successful. In addition, our repeated requests to BD to bring us into contact with the relevant technicians went unanswered. Consequently, we were forced to restrict our quantitative analysis to cells whose GFP levels were at least twice the autofluorescence level (corresponding the log-mean levels roughly equal to those of the dimmest calibration beads in Suppl. Fig. S4).

### 1.2 Mixture model of the scatter measurements

The scatter of each measured event is characterized by 4 statistics: the height and width of both the forward- and side-scatter. We fit the set of 4D scatter measurements for a given dataset by a mixture of a multivariate Gaussian and a uniform distribution. The idea is that the Gaussian distribution models the viable cells, while the uniform distribution represents outliers that may correspond to fragments of dead cells or other debris. Concretely, we assume that the probability to observe the 4D vector 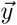 for a single measured event is given by

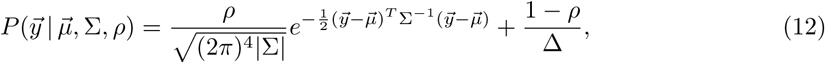

where 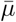 is the center of the 4D multivariate Gaussian, Σ its covariance matrix, *ρ* the fraction of viable cell measurements in the dataset, and ∆ is the volume of the 4D hypercube spanned by all the data points. Note that |Σ| denotes the determinant of the covariance matrix. The *E-Flow* package that we distribute contains a C++ implementation where the model (12) is fit to a given dataset using expectation maximization. An example of the results of this fitting to a dataset of *E. coli* cells grown in minimal media with lactose is shown in Fig. 2 of the main paper, showing that the observed 4D scatter can be well approximated by the mixture model.

Once the mixture model is fitted, we can calculate a posterior probability *p*_*i*_ for each observation *i* that it derives from the Gaussian mixture component, i.e. that it is a viable cell measurement. By default *E-Flow* filters out all events *i* for which this posterior probability is less than 0.5 and retains all measurements with posterior probability larger than 0.5, but users can choose to alter this threshold in posterior probability. Supplementary Figure S5 shows the same 4D scatter of measurements as shown in Fig. 2 of the main text, but now with all selected events in red, and all events that were filtered out in black.

To investigate the effect of the filtering strategy on the observed fluorescence levels we calculated the distribution of fluorescence levels of several reporters when using either a very strict threshold, retaining only cells with posterior *p >* 1 − *ϵ*^−10^ or a very lenient threshold, retaining all cells with *p > e*^−10^. As shown in Suppl. Fig. S6 there is virtually no difference between the observed distribution of fluorescences with the two threshold. This observation strongly suggests that it is not possible to effectively pick out a subpopulation of cells of similar size by strict gating on forward- and side-scatter.

### 1.3 Error of forward- and side-scatter as a measure of cell size

As explained in the main text, we used the known accuracy of microscopy measurements to estimate the accuracy of forward- and side-scatter as a measure of cell size. Suppl Fig. S7 shows the standard-deviations of the cell size estimates using either the height or width of either the forward- or side-scatter as a function of the total standard-deviation in the signal.

The figure shows that the width of forward- and side-scatter signals are completely uninformative for cell size, i.e. they lie on the line *y* = *x* where the variance in the measured signal var(*x*_*m*_) is equal to the measurement noise var(*ϵ*). Although the height of the forward- and side-scatter signals carry some information about cell size, the measurement noise is still larger than the true variability in cell size, so that these measures are too noisy to be effectively used as measures of cell size. In contrast, microscopy measurements have only 3% errors whereas true size varies by about 20% across cells.

#### 1.3.1 Inferring the mean and variance of the distributions of fluorescence levels

After filtering events based on forward- and side-scatter, we next fit the distribution of fluorescence by a mixture of a log-normal distribution, which captures valid measurements and corresponds to the bulk of the measurements, and a uniform distribution capturing outliers. In particular, we assume that the probability for a fluorescence measurement to have log-fluorescence *x* is given by the mixture:

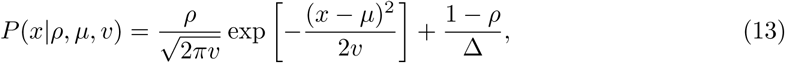

where *µ* and *υ* are the mean and variance of log-fluorescence levels, *ρ* is the fraction of ‘valid’ measurements and ∆ = *x*_max_ − *x*_mix_ is the observed range of log-fluorescence levels. The log-likelihood for a dataset of *n* measurements *x*_*i*_ is then simply given by

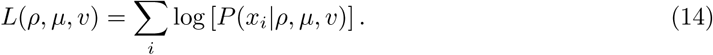

We maximize the log-likelihood (14) using expectation maximization to obtain the fitted parameters *ρ*_∗_, *µ*_∗_ and *υ*_∗_. To obtain error bars *σ*_*µ*_ and *σ*_*υ*_ on the mean *µ* and variance *υ* we obtain the Hessian matrix of the log-likelihood by expanding to second order around its maximum and we take the diagonal elements of its inverse. For any given set of fluorescence measurements, our package *E-Flow* returns both the fitted values (*µ*_∗_, *υ*_∗_) and their error bars (*σ*_*µ*_, *σ*_*υ*_).

Below we develop methods for correcting the observed fluorescence levels for both cell autofluorescence and for the shot noise in the FCM measurements. In order to do this we also need the mean and variance of fluorescence levels, rather than just the mean and variance of *log*-fluorescence levels. That is, if we write for the fluorescence of a cell *f* = *e*^*x*^, we need to obtain the mean and variance of *f*. Using the well-known expressions of the mean and variance of a log-normal distribution we have

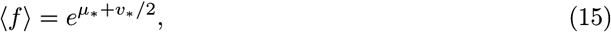

and

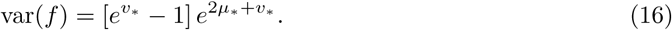

Further, given that the error bars on the mean and variance are generally small compared to their means, we use a linear approximation to calculate error bars on the mean and variance of *f*, and find

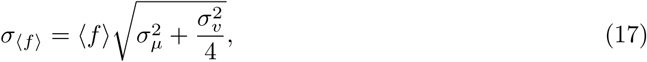

and

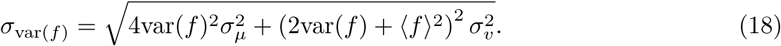

The *E-Flow* package also returns these estimates and error bars for fluorescence levels *f*.

### 1.4 Averaging of replicate autofluorescence measurements

As shown in Fig. 4 in the main text, we measured the autofluorescence distributions of cells that do not express GFP in triplicate on 4 separate days, and for two different strains. Noticing that there appears to be a small but systematic variation in both the means and variances of the autofluorescence levels, we proceeded as follows to estimate an overall average for the mean and variance of autofluorescence, adapting a method first presented in [13].

We assume that there is an overall mean autofluorescence *µ* and that on any given day *d*, the mean autofluorescence *µ*_*d*_ on that day deviates from *µ* by some amount *δ*_*d*_, i.e.

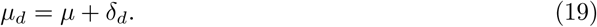

We will assume that the deviation *δ*_*d*_ varies by an (unknown) amount *τ* across days, following a Gaussian distribution. That is, the probability of having mean autofluorescence *µ*_*d*_ on any given day is

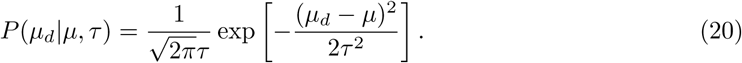

Let *µ*_*id*_ be the estimated mean autofluorescence of replicate *i* on day *d*, and let *σ*_*id*_ be the associated error-bar on this mean. Assuming that the true mean autofluorescence on day *d* was *µ*_*d*_, the probability to obtain a measured mean autofluorescence *µ*_*id*_ is also given by a Gaussian distribution:

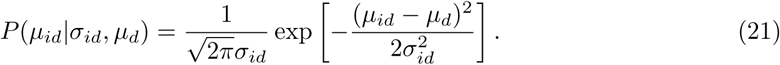

The probability of the data *D*, i.e. all replicate measurements *µ*_*id*_ given an overall mean *µ* and variance across days *τ* ^2^ is given by taking the product over all measurements and days, and marginalizing over all day-dependent means *µ*_*d*_:

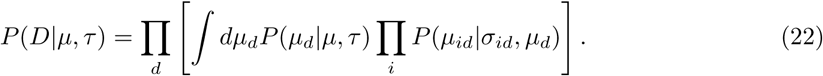

These integrals can all be performed analytically and we obtain

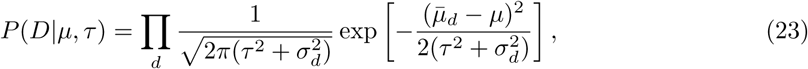

where we have defined the measured mean 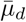 on day *d* as

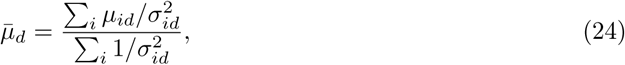

and the squared error 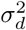 on the measured mean on day *d* as

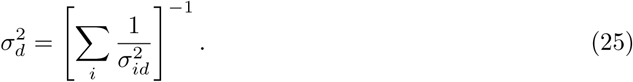

If we define the weighted overall average

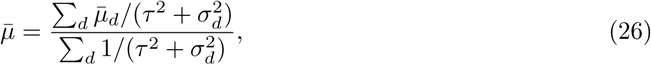

and the overall squared error

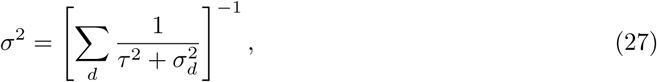

then we can rewrite equation (23) as

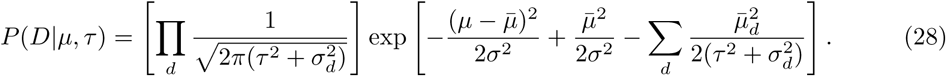

We can then finally marginalize over *µ* to obtain a likelihood that depends only on *τ* and the measurements:

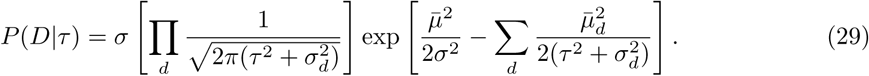

We maximize equation (29) with respect to *τ* to find the optimal value *τ*_∗_. We then substitute this value *τ*_∗_ into equations (26) and (27) to obtain a final estimate 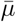 of the overall mean autofluorescence and an error bar *σ* on this estimate. The same procedure is used to estimate average variances 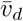 for each day and an overall average variance 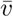. The *E-Flow* package returns all these values as the overall estimates of autofluorescence averaged over multiple replicates and days.

### 1.5 Fitting the shot noise strength using calibration beads

The electronic cascade in the photomultiplier tube introduces a noise whose variance can be described as a Gaussian with a variance proportional to the square root of the incoming signal [11]. This means that, for every measurement of fluorescence, the relationship between measured intensity *I*_*M*_ and true intensity *I*_*T*_ is given by

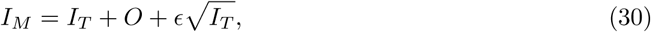

with 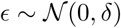 and *O* is an offset introduced by the electronics in order to avoid negative values when *I*_*T*_ is very small.

To infer the strength *δ* of the shot noise term we used a set of calibration beads consisting of a mixture of beads of three different expression levels. The top right panel of Suppl. Fig. S8 shows the observed distribution of expression levels for one set of beads showing three main peaks and one small additional peak at very high intensity. We believe that this highest peak is an artefact due to aggregation of multiple beads. We fit the distribution of bead intensities to a mixture of a uniform distribution (to account for outliers) plus four Gaussians. We discard all points assigned to the uniform distribution and the highest Gaussian component, and estimated the means, variances, and *CV*^2^ for each of the remaining three Gaussians. This procedure was repeated for multiple datasets. In total we estimated means, variances, and *CV*^2^ for 31 datasets of calibration beads. The top left panel of Suppl. Fig S8 shows the *CV*^2^ as a function of the mean intensities of the beads across all datasets showing that *CV*^2^ drops approximately inversely with mean expression, as expected for shot noise.

Given one dataset of beads expressions, we can now fit a model based on Eq. (30). First of all it can be proved that under this model the mean and variance of the expression intensities of one single peak *i* are given by

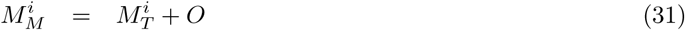

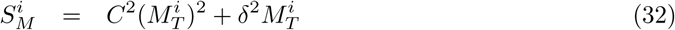

where we have supposed that the true oefficient of variation *C* is the same for every peak (we assume that each of the 3 types of beads has the same manufacturing accuracy in their fluorescence intensities). From the histogram in Fig. S8 (top right) it is clear that in logarithmic space the intensities are well described by Gaussian distributions, which means that 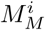 and 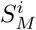 are the mean and variance of a log-normal distribution. We can work out the *µ* and *σ* of a normal distribution in log-space that would have 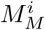 and 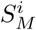 as mean and variance in real space

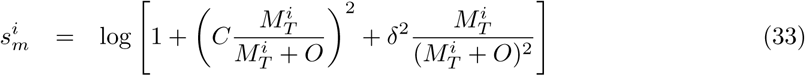

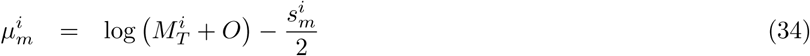

It follows that in logarithmic space the probability of observing a data point *x*_*k*_ (including also a uniform distribution that takes care of possible outliers) is

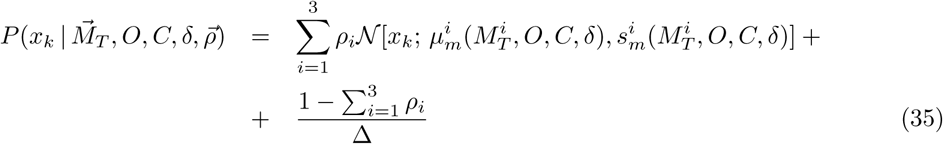

where 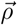 is a vector of weights whose elements sum to one and ∆ is the range of the data in logarithmic space. The log-likelihood of the data is

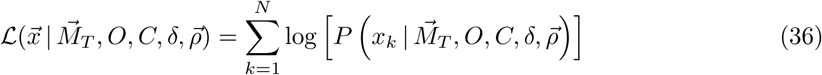

This likelihood is maximized using a coordinate descent algorithm and the error bar on *δ* was estimated by expanding the log-likelihood to second order around its maximum, keeping all other parameters fixed. The same procedure was used to estimate an error bar on the offset *O*.

After obtaining estimates of *δ* and *O* and their error bars for each dataset, we averaged the *δ*s and *O*s from different datasets in the same way as we did for autofluorescence measurements above. The bottom part of Suppl. Fig. S8 shows the estimates of the shot noise and offset for different datasets and the final average of all these estimates. We find that *δ* has a value of 12.7 with a variation of 0.6 among datasets. Finally, the black line on the top left panel of Suppl. Fig. S8 shows that the model correctly describes the observed decrease in *CV*^2^ with mean.

### 1.6 Mother machine statistics

#### 1.6.1 Age distribution in the Mother machine

Especially for highly expressed genes the GFP concentration varies relatively little across cells, so that the total amount of GFP in a cell correlates well with its size. Since each cell expands exponentially along its division cycle in exponential growth, cell size correlates well with the ‘age’ of a cell, i.e. where it is in its division cycle. For this reason we expect a correlation between the total GFP distribution and the age distribution of the population, i.e. the relative frequencies of cells in different stages of their division cycle. Thus, in order to meaningfully compare GFP distributions from the FCM and from cells growing in a microfluidic device, we must check that the age distributions of the two populations are indeed comparable.

The cell population analyzed in the FCM is expected to have an over-representation of younger cells, since for every old cell that divides two young cells are produced whereas, in the microfluidic device, we expect less over-representation of young cells due to the fact that cells continuously leave the growth channel.

To infer the population structure of a growing population of cells, we use the Leslie approach [14]. We discretize the time in steps of 3 minutes, corresponding to the acquisition time of the time lapse microscopy, and we look for the probability *P* (*a, t*) of having cells of age *a* at time *t*. The Leslie theory is based on two quantities, the fecundity *f* (*x, t*) which represents the expected contribution from an individual aged *x* at time point *t* to the population aged 1 at time point *t* + 1, and the probability *P* (*x, t*) of survival over one time point for individuals who are present in age-class *x* at time *t*. These quantities form the Leslie matrix, which is defined as

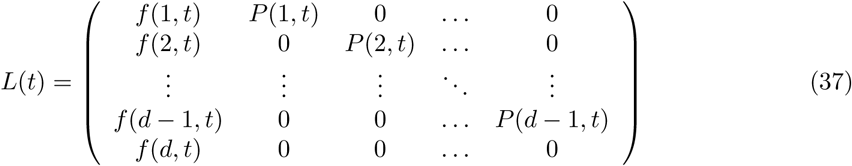

this matrix is a squared *d × d* matrix where *d* is an upper limit on the age of the cells, after which *P* (*x, t*) = 0. The population structure at time *t* is given by a vector 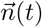 of dimension *d*, where the entry *i* is the number of individual in the population of age *i*. The theory states that

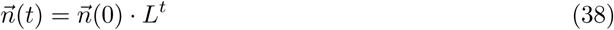

where we use the convention that vectors are represented by rows. If the matrix *L* is diagonalizable and its entries are independent of time, it can be shown that for very large *t* the population reaches a stable age distribution with

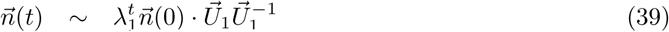

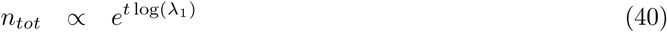

where *λ*_1_ and 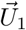 are the leading eigenvalues and eigenvectors of *L*.

From the data from the microfluidic device we can estimate the probability for a cell of age *x* at time *t* to surive one more time point by computing the probability that the division occurs at an age larger than *x*; the fecundity is then given as twice the probability that the cell doesn’t survive. Suppl. Fig. S9 shows the computed *P* (*x, t*) by considering cells at different times *t* from the beginning of the experiment and it shows that they are all the same, suggesting that the Leslie matrix is indeed independent of time. We thus decided to compute a single curve for *P* (*x*) by pooling all the cells together independently of the time at which they were measured. The resulting curve shows that the survival probability becomes negligible after 150 minutes, with a medium age at division of 80 minutes.

We found that the exponential growth rate of the population, Eq (40), is *ρ* ~ 0.0089/*min*, which is the same as the measured single cell growth rate. As expected, the population age structure differs markedly between the observed and the theoretical one, as shown on the bottom right part of Fig S9. In particular the theoretical distribution predicts more young cells than old, while in the microfluidic device we have a nearly uniform age distribution with a right tail; hence, as expected, young cells are relatively under-represented in the population in the microfluidic device.

Assuming that the GFP concentration is constant, the total GFP must depend on the cell age through the length and in computing the mean and variances we have to weight the GFP by the ratio between the theoretical and the observed age fractions. Specifically, if *P*_*obs*_(age) and *P*_*th*_(age) are the observed and the theoretical age distributions, then for the log(*GFP*) distribution it holds

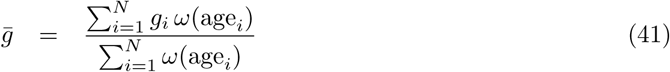

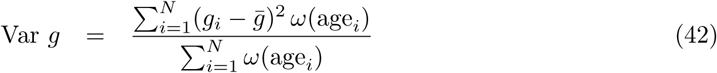

where *i* runs over all cell observations, *g* ≡ log(*G*) and *ω*(age) is a weighting factor that accounts for the over-representation of older cells

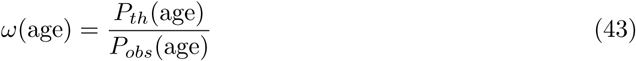

Going into log-space, we have estimates more robust against outliers and we can approximate the *CV*^2^ in the real space as the variance in the log-space. Suppl. Fig. S10 shows that the age structure correction makes the CVs only slightly lower than the one measured ignoring the age structure effects.

#### 1.6.2 Correlation in the measurements

As we are recording the size and GFP expression of cells every 3 minutes in the microfluidic device, observations from nearby time points are clearly correlated, and it could be suspected that this may affect the statistics we calculate. In contrast, in the FCM, we have cells measured at only one time point, so we don’t have correlations coming from the fact that we have multiple data points coming from the same cell. To check whether the presence of these correlated data points can substantially bias the statistics, we measured the *CV*^2^ for different promoters and media on different dates in two different ways:

1. Taking all the data, regardless of the fact that they may come from the same cell.
2. Taking only one randomly selected data point for each cell.

The first condition is the one that we use throughout the paper to compute the statistics, while the second one should be closer to the FCM setup. Suppl. Fig. S10 shows that the *CV*^2^ are almost identical whether correlated time points are included or not. In conclusion, neither the age structure nor the correlated measurements affect the comparison of *CV*^2^ between measurements from the FCM and from the microfluidic device.

**Fig S1.**
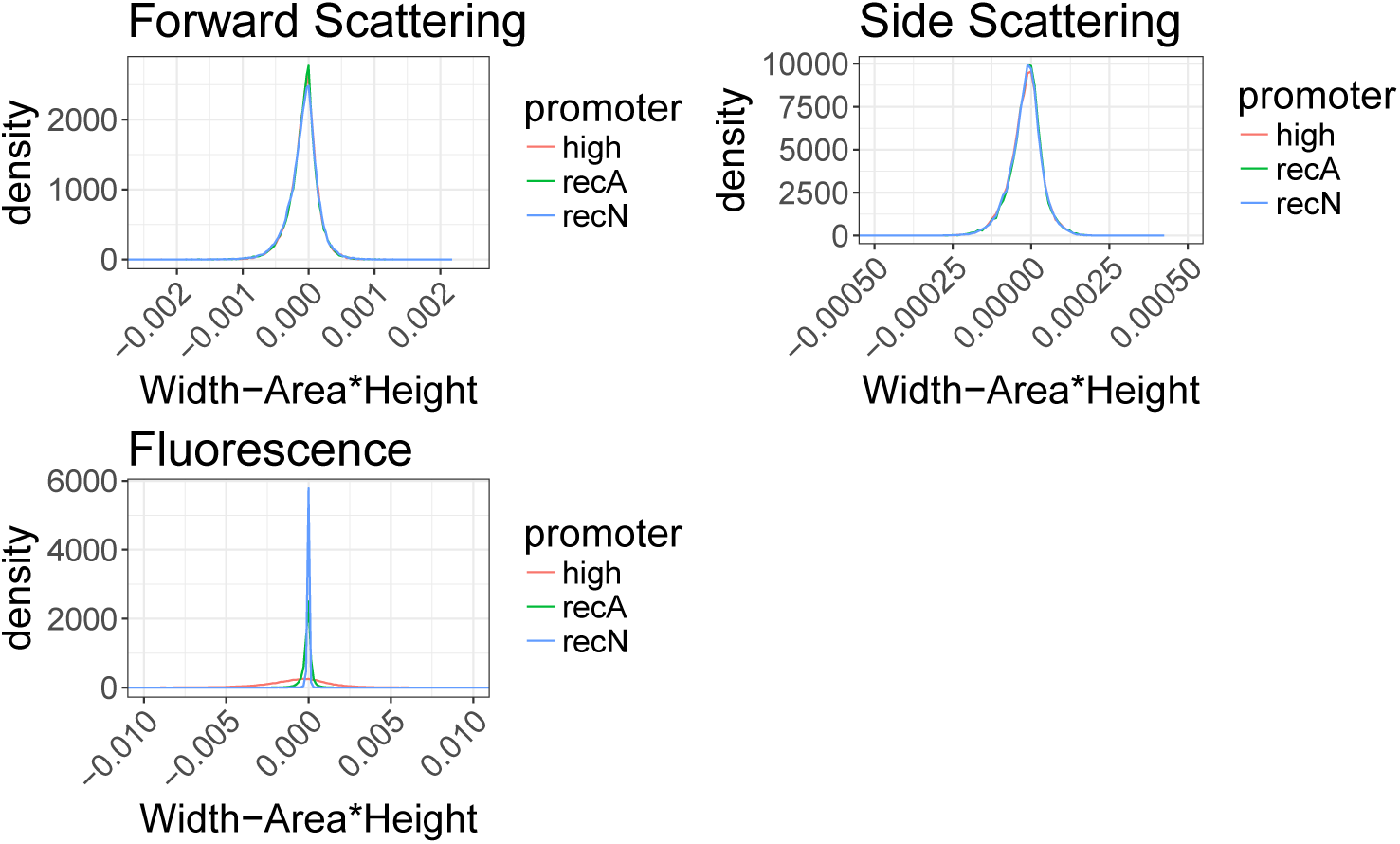
Pulse area is proportional to the product of height and width. Apart from very rarely occurring negative or saturated signals, the reported area of each pulse us equal to a constant *C* times the product of the reported height and width. The histograms show the distribution of area – *C* ⋅ width ⋅ height, which is highly concentrated around zero. Different colors represent different data-sets acquired on different days using different promoters.

**Fig S2.**
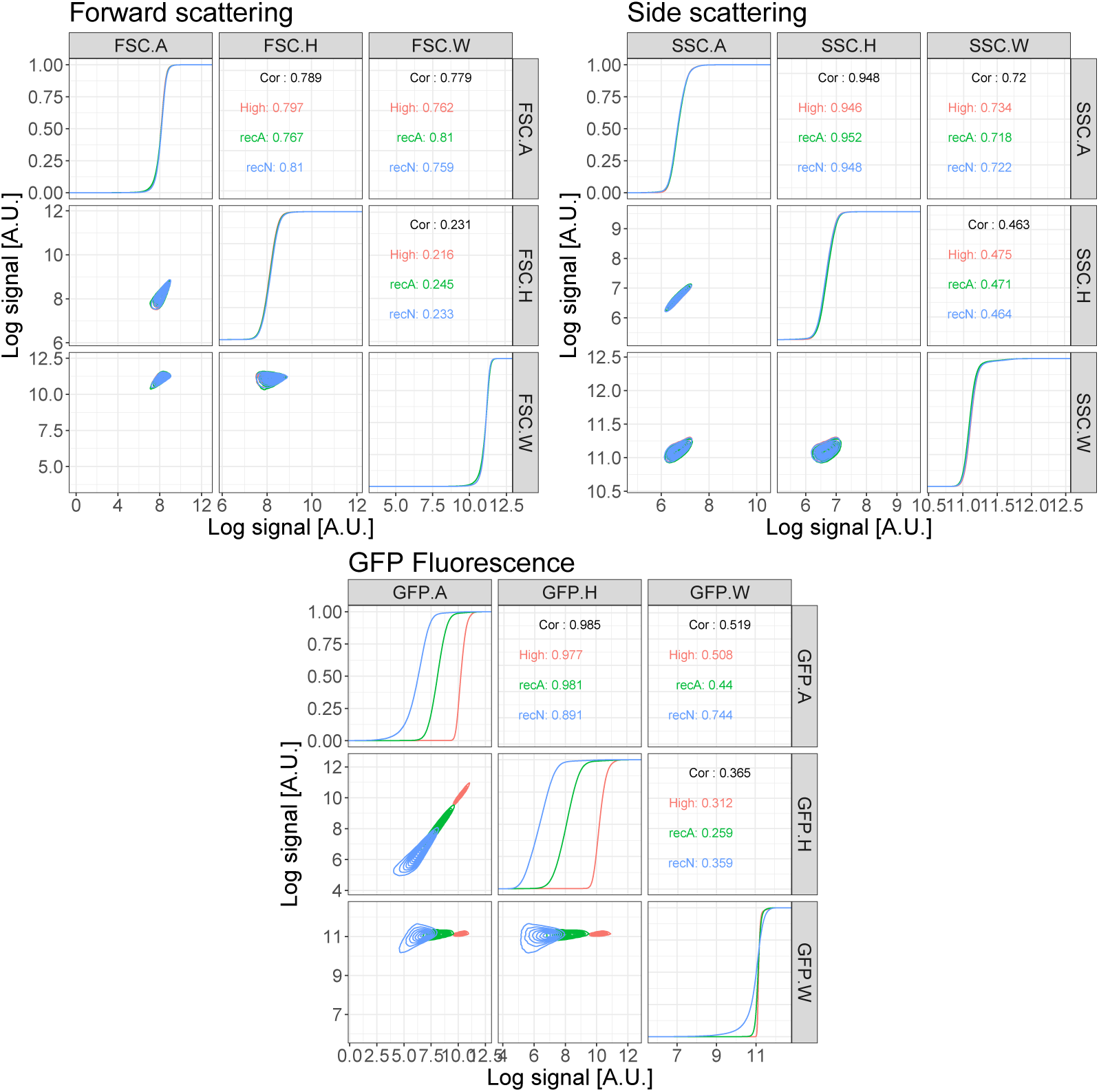
Correlations among the different statistics reported for each signal pulse. The plots show the correlations among the height, area and width of the signal pulses for the forward- and side-scatter as well as the fluorescence measurements. The colors represent 3 different promoters, two native (recA and recN) and one synthetic (high).

**Fig S3.**
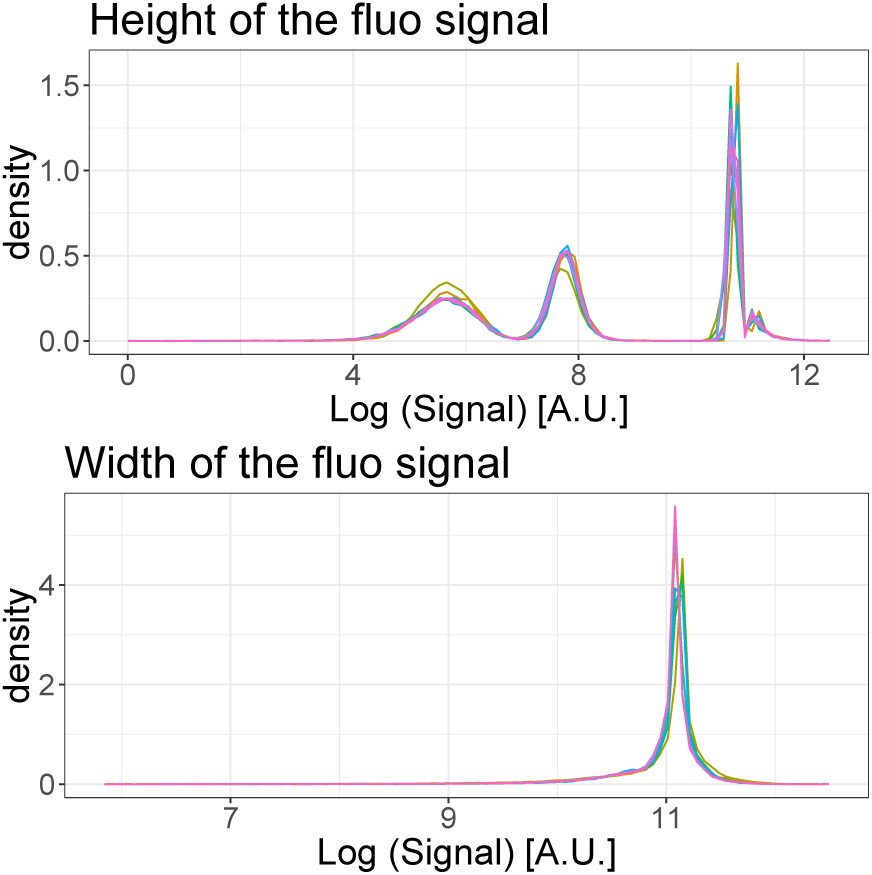
Population separation of calibration beads based on their scattering profile. Distributions of the measured heights (top) and widths (bottom) of the fluorescence signal pulses for populations of artificial beads. Each color corresponds to one measurement run of a set of artificial beads consisting of a mixture of beads of three different fluorescent intensities. We see that, whereas signal height shows 3 clearly separated peaks, the distribution of signal widths is narrowly peaked at a single value. Note also the notably wider variance of the peaks at low intensities than at high intensities which results from measurement shot noise.

**Fig S4.**
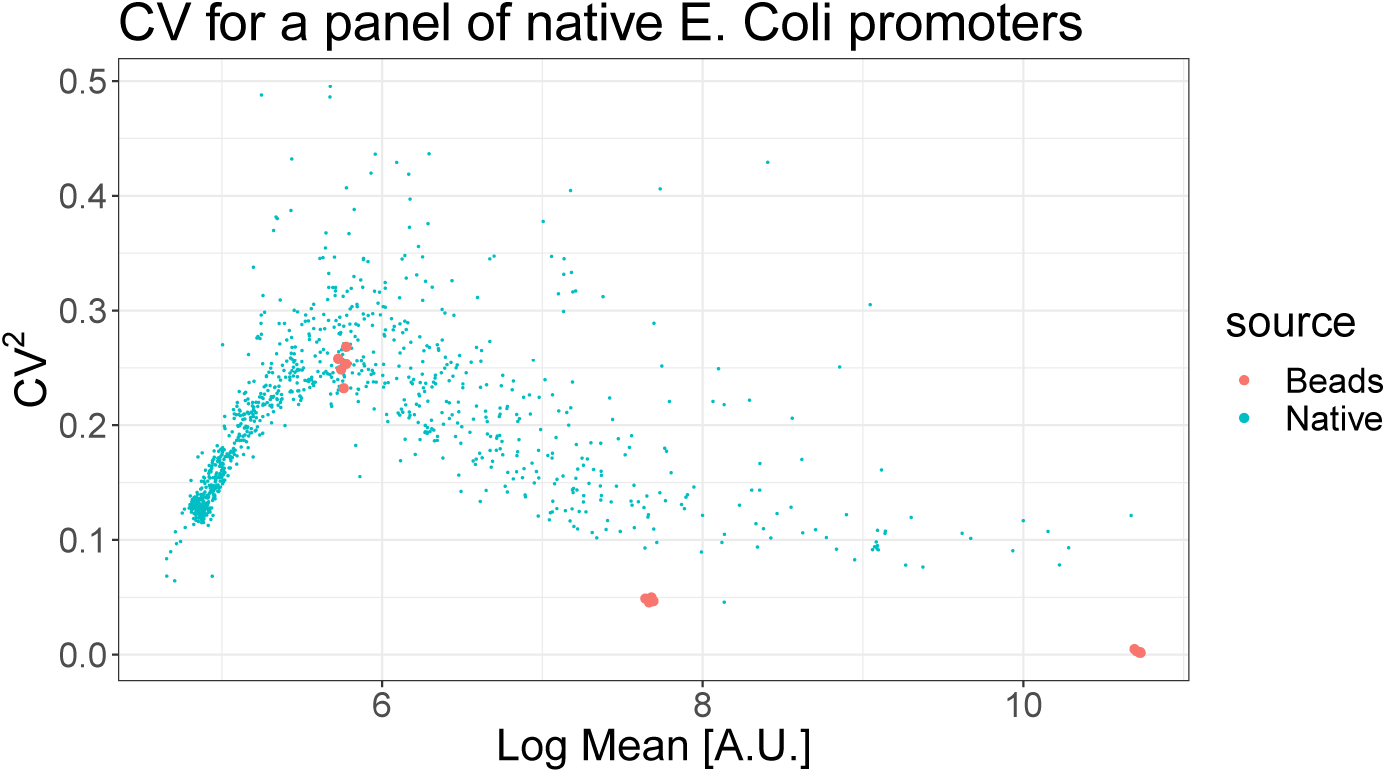
Coefficient of variation as a function of mean fluorescence level exhibits anomalous behavior at low fluorescence. The blue scatter shows the measured coefficient of variation squared *CV*^2^ (vertical axis) as a function of the logarithm of mean fluorescence level (horizontal axis) for a library of transcriptional reporters of native *E. coli* promoters [5], with each blue point corresponding to one promoter of the library. The red points show the same statistics for the calibration beads at three intensities. Note that, due to shot noise in the fluorescence measurements, the *CV*^2^ increases as the mean fluorescence decreases, including for the low-variance calibration beads. However, at very low fluorescence, the *CV*^2^ reaches a maximum and decreases for even lower fluorescence.

**Fig S5.**
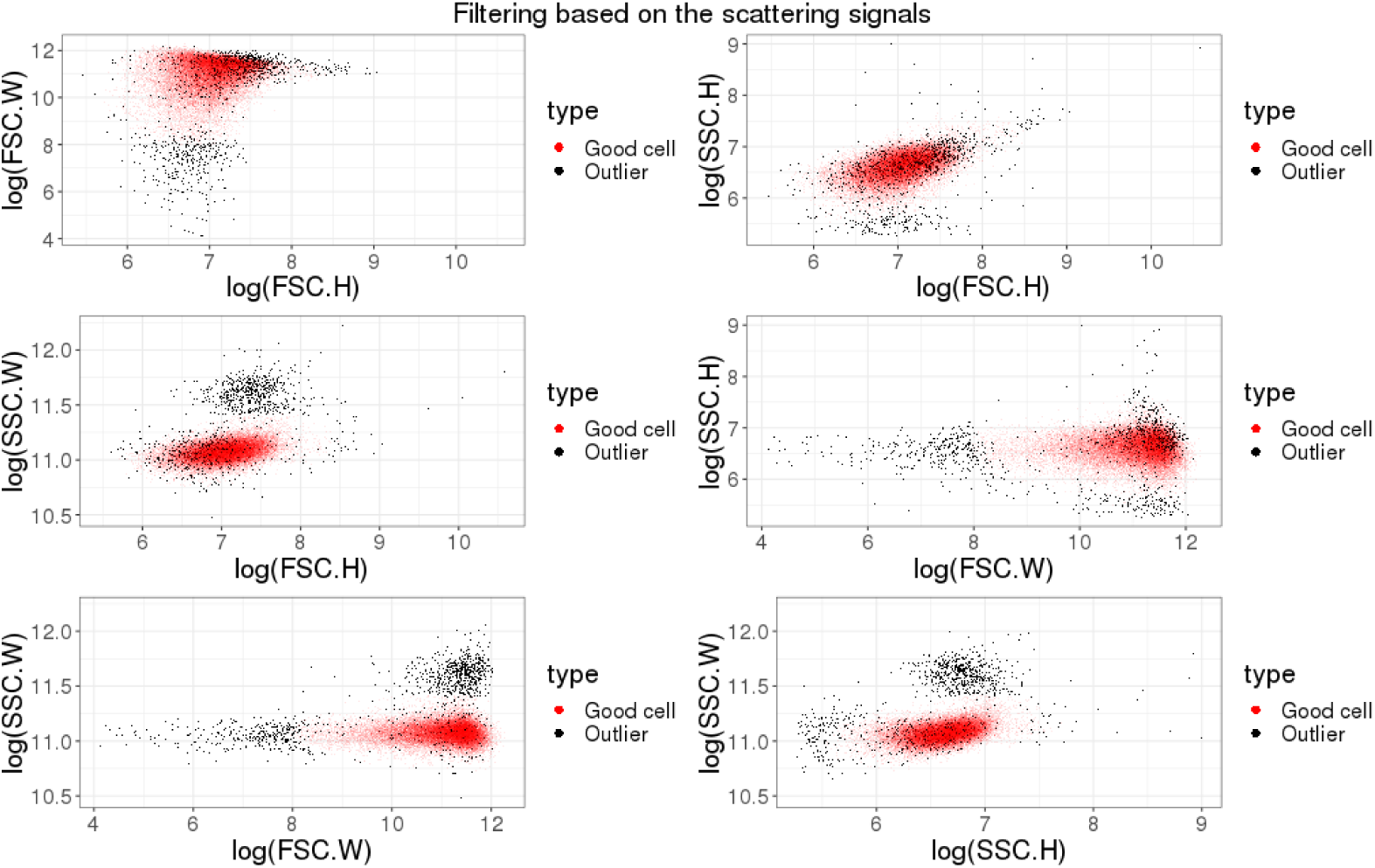
Filtering of events based on their scattering profiles. The panels show a scatter of the forward- and side-scatter measurements for a set of *E. coli* cells grown in minimal media with lactose. Events that have posterior probability larger than 0.5 to stem from the Gaussian component of the mixture are indicated in red, and events with posterior less than 0.5 are superimposed in black. The red events are considered viable cell measurements and the black events are discarded. Note that around 4% of the measured events are discarded for this dataset.

**Fig S6.**
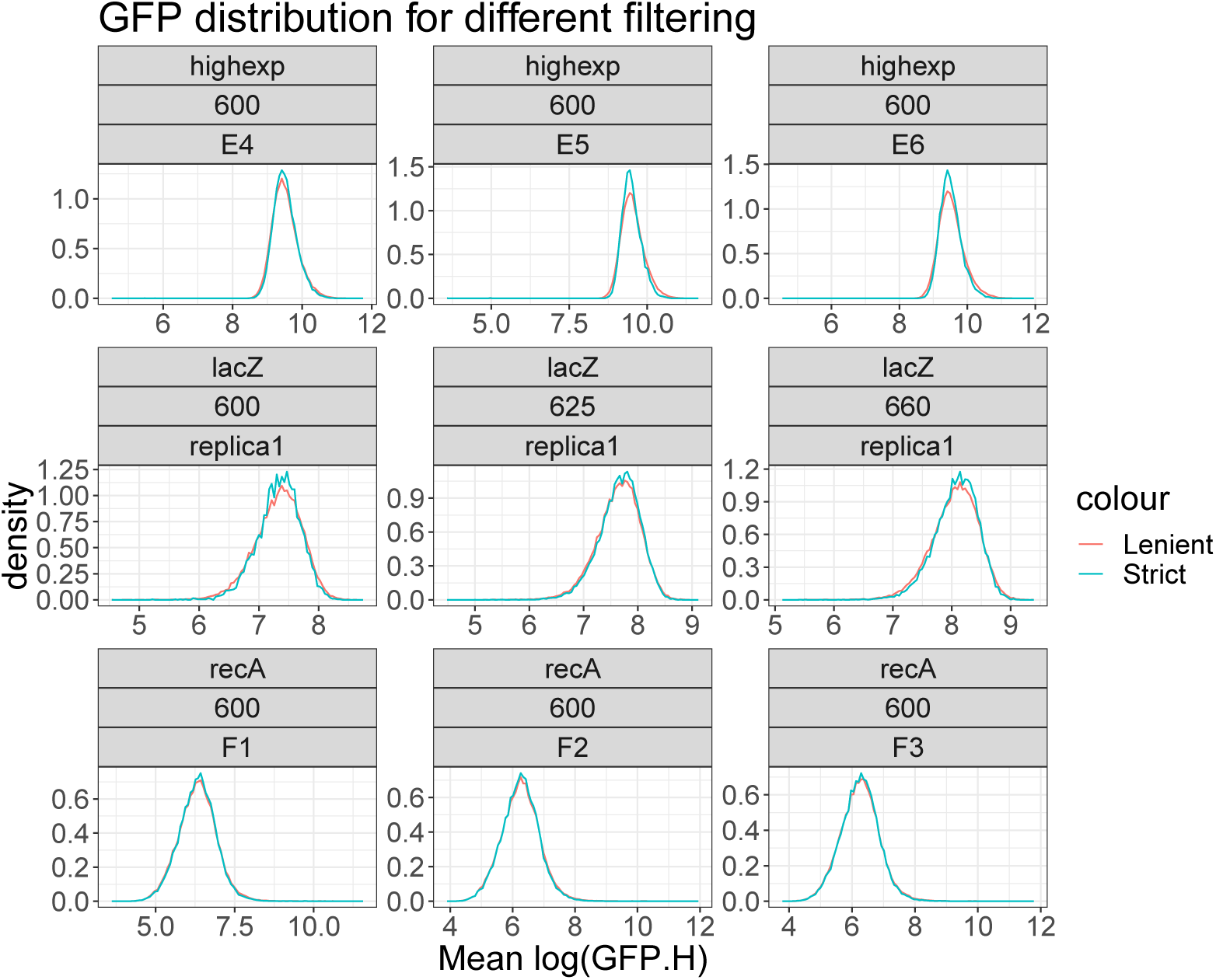
Fluorescence distribution when using very strict or very lenient gating of forward- and side-scatter. Each panel shows the observed distribution of fluorescence levels of a transcriptional reporter (rows) using different replicate measurements (columns) when using either a very strict threshold *p >* 1 − *ϵ*^−10^ or a very lenient threshold *p > e*^−10^ for filtering events. Note that the name of the reporter and the voltage of the flow cytometer’s laser used in a given replicate are indicated at the top of each panel.

**Fig S7.**
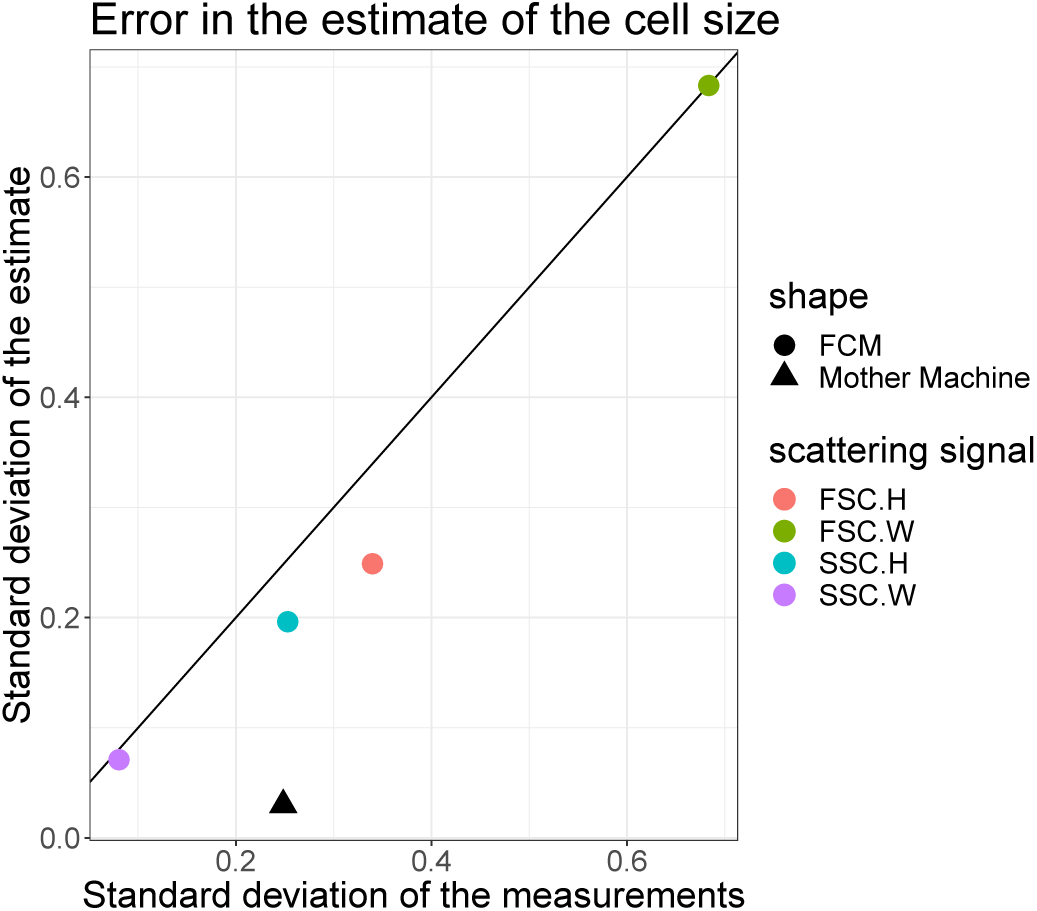
Estimated errors of forward and side-scatter as measures of cell size. Using a comparison of measured variances of microscope and FCM measurements on the same cells in the same conditions, we estimated the errors of forward- and side-scatter as measures of cell size (see main text). The plot shows the estimated standard-deviations of cell size estimates 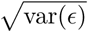 using either height or width of forward- or side-scatter measurement in the FCM (colored discs) as a function of the total standard-deviation in the measurements 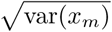. For comparison, the black triangle shows the standard-deviation of the microscopy cell size measurements. The black line shows the function *y* = *x* corresponding to measurements that are completely uninformative, i.e. dominated by measurement noise.

**Fig S8.**
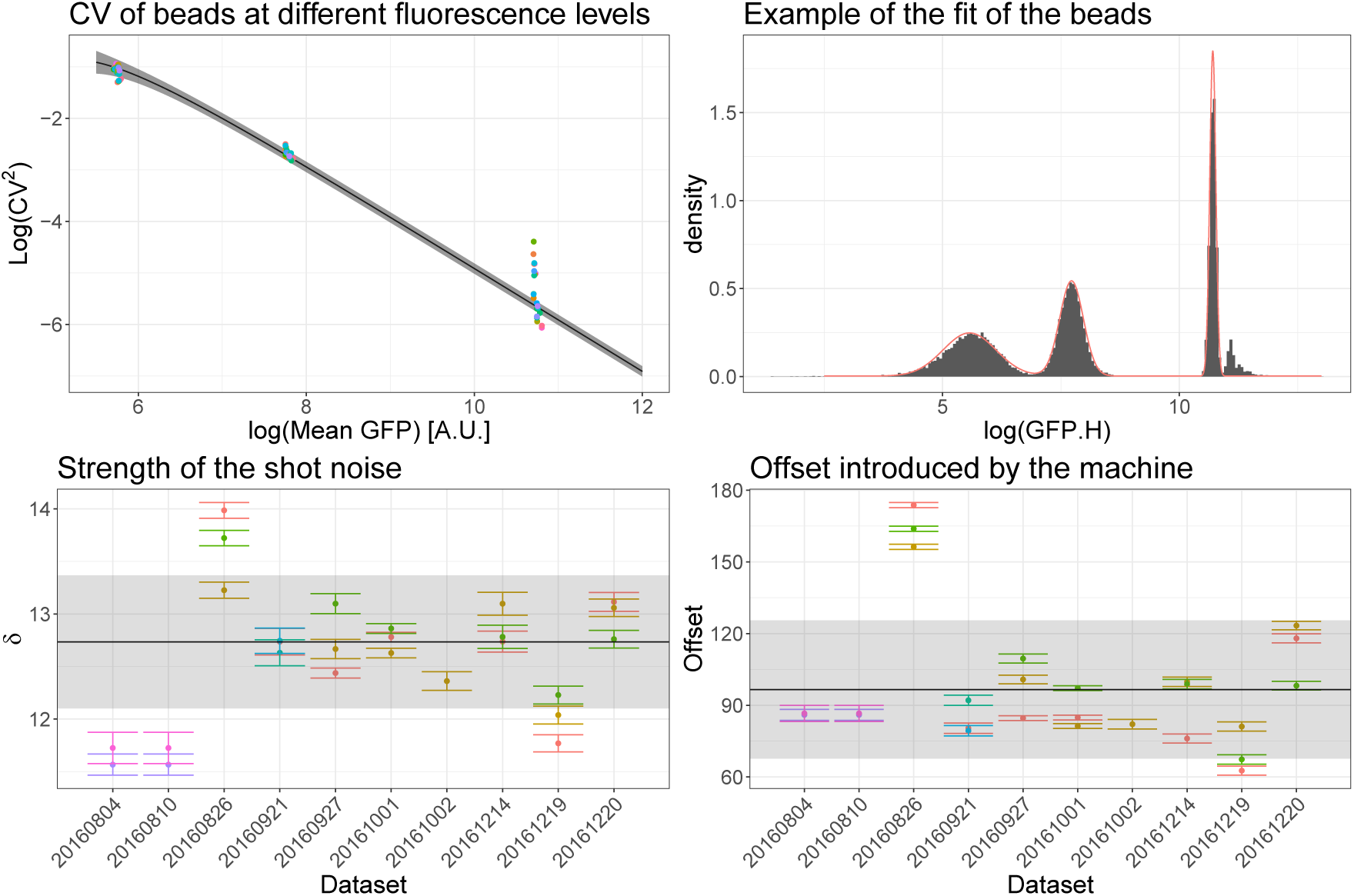
Estimation of the shot noise strength using calibration beads. Top left: *CV*^2^ as a function of mean intensity for each Gaussian component of each mixture of beads analyzed. Both axes are shown on a log-scale and different colors refer to different datasets. The black line and the shaded region represent the fit obtained using a shot noise strength *δ* = 12.7 ± 0.6. Top right: Example of the intensity distribution of a single dataset of beads; the red line shows the fit with a mixture of three Gaussians and one uniform. Bottom: Estimates of the strength *δ* (left) and offset *O* (right) for different datasets; the estimation is done fitting a model of the form given by equations (35) and (36). The black lines correspond to the means of *δ* and *O* and the gray regions correspond to their variation among datasets.

**Fig S9.**
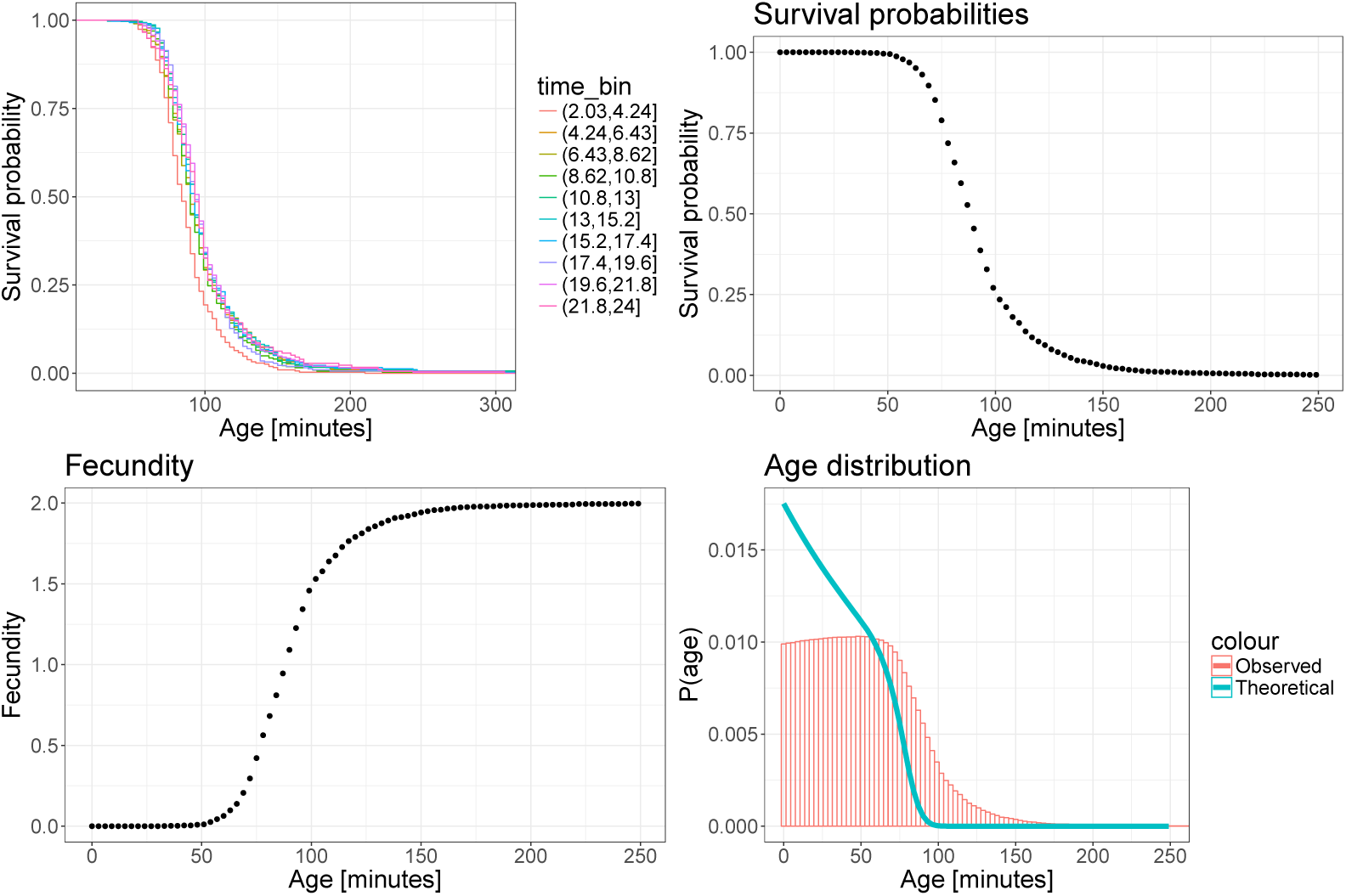
The Leslie model. *Top left*: The one-time point survival probabilities don’t depend on the time and they show that the survival probability becomes negligible for cells older than about 150 minutes. The medium age at division is around 80 minutes. *Top right*: Since the probabilities don’t depend on time, we pooled data from all times together to obtain a single survival curve. *Bottom left*: Fecundity as function of cell age. *Bottom right* Asymptotic age distribution of the population for large times as obserced (red) and predicted by the theory (blue). Note that the observed distribution has an over-representation of old cells relative to the theoretically predicted distribution.

**Fig S10.**
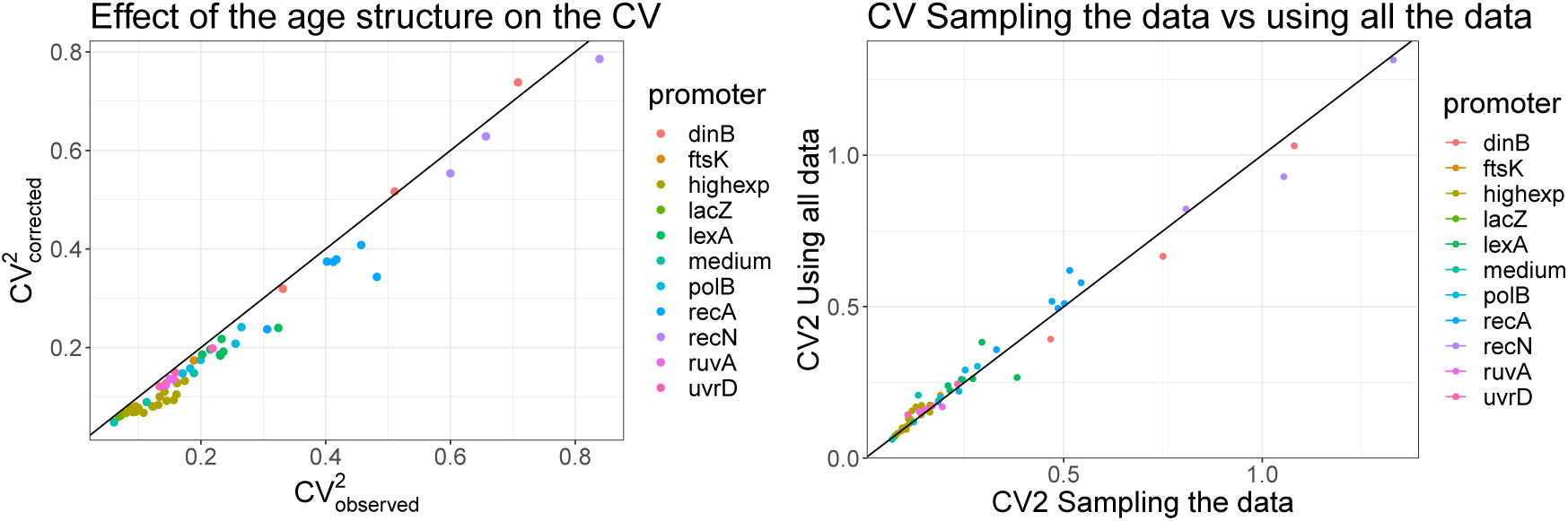
Effect of the age structure on the measured *CV*^2^. *Left*: *CV*^2^ corrected by the age structure in the microfluidic device versus the *CV*^2^ as directly measured, ignoring the age structure effect. The age structure has the effect of making the measured *CV*^2^ slightly higher than the *CV*^2^ corrected by the age structure of the population. *Right*: *CV*^2^ obtained by taking all the measures in the microfluidc device versus the *CV*^2^ when taking only one point, chosen at random, per cell. We can see that the difference is very small and the points accumulate around the bisector.

